# Empirically establishing drug exposure records directly from untargeted metabolomics data

**DOI:** 10.1101/2024.10.07.617109

**Authors:** Haoqi Nina Zhao, Kine Eide Kvitne, Corinna Brungs, Siddharth Mohan, Vincent Charron-Lamoureux, Wout Bittremieux, Runbang Tang, Robin Schmid, Santosh Lamichhane, Yasin El Abiead, Mohammadsobhan S. Andalibi, Helena Mannochio-Russo, Madison Ambre, Nicole E. Avalon, MacKenzie Bryant, Andrés Mauricio Caraballo-Rodríguez, Martin Casas Maya, Loryn Chin, Ronald J. Ellis, Donald Franklin, Sagan Girod, Paulo Wender P Gomes, Lauren Hansen, Robert Heaton, Jennifer E. Iudicello, Alan K. Jarmusch, Lora Khatib, Scott Letendre, Sarolt Magyari, Daniel McDonald, Ipsita Mohanty, Andrés Cumsille, David J. Moore, Prajit Rajkumar, Dylan H. Ross, Harshada Sapre, Mohammad Reza Zare Shahneh, Sydney P. Thomas, Caitlin Tribelhorn, Helena M. Tubb, Corinn Walker, Crystal X. Wang, Shipei Xing, Jasmine Zemlin, Simone Zuffa, David S. Wishart, Rima Kaddurah-Daouk, Mingxun Wang, Manuela Raffatellu, Karsten Zengler, Tomáš Pluskal, Libin Xu, Rob Knight, Shirley M. Tsunoda, Pieter C. Dorrestein

## Abstract

Despite extensive efforts, extracting information on medication exposure from clinical records remains challenging. To complement this approach, we developed the tandem mass spectrometry (MS/MS) based GNPS Drug Library. This resource integrates MS/MS data for drugs and their metabolites/analogs with controlled vocabularies on exposure sources, pharmacologic classes, therapeutic indications, and mechanisms of action. It enables direct analysis of drug exposure and metabolism from untargeted metabolomics data independent of clinical records. Our library facilitates stratification of individuals in clinical studies based on the empirically detected medications, exemplified by drug-dependent microbiota-derived *N*-acyl lipid changes in a cohort with human immunodeficiency virus. The GNPS Drug Library holds potential for broader applications in drug discovery and precision medicine.

## Main Text

Growing evidence suggests that the chemical exposome plays a critical role in shaping human health, with drugs being a significant source of chemical exposure that carries profound health implications.^1^ According to a recent survey by the Center for Disease Control and Prevention, nearly half (45.7%) of the U.S. population used at least one prescription drug in the past 30 days.^2^ Drug concentrations in human blood are on par with those of endogenous and dietary molecules,^3^ and have important impacts on the metabolic states and microbiome composition.^4–7^ Clinical research typically relies on medical records or self-reporting surveys to assess drug exposure,^8^ but these methods are costly and often incomplete.^8–10^ They often overlook over-the-counter medications and supplements, and fail to account for patient adherence. Additionally, they miss drug usage not documented in medical records, such as those purchased online,^10,11^ acquired across borders,^12,13^ or consumed through secondary use of leftover drugs. Medical records are also incapable of documenting drugs introduced into the food supply that are unknowingly consumed, such as the antifungal natamycin used both to treat fungal eye infections and as a preservative for dairy products. Additionally, the varying half-lives of drugs and their metabolites further complicate exposure assessment, as some drugs are rapidly eliminated from the body while others can persist for months.^14,15^

Untargeted metabolomics offers the opportunity to complement clinical records by empirically establishing the presence of drugs and their metabolites directly from biological samples. However, liquid chromatography-tandem mass spectrometry (LC-MS/MS) based annotations, which rely on reference MS/MS library matches, are difficult to interpret. For example, annotation may return a complex IUPAC chemical name like “(2R,3S,4R,5R,8R,10R,11R,12S,13S,14R)-11-[(2S,3R,4S,6R)-4-(dimethylamino)-3-hydroxy-6-methyloxan-2-yl]oxy-2-ethyl-3,4,10-trihydroxy-13-[(2R,4R,5S,6S)-5-hydroxy-4-methoxy-4,6-dimethyloxan-2-yl]oxy-3,5,6,8,10,12,14-heptamethyl-1-oxa-6-azacyclopentadecan-15-one”. A text search in the right reference resource or an open web search can hopefully link this IUPAC name to “azithromycin”, the drug name used in clinical settings. A second search of the term “azithromycin” is then required to connect the name to its therapeutic role, in this case an antibiotic originally isolated from a bacterium. While this example involves a simple name and a limited number of identifiers for azithromycin, other compounds, like penicillin G or aspirin, have hundreds of synonyms and identifiers in chemistry databases such as PubChem, making the identification process more challenging. This task must be repeated for every obtained annotation, which can range from hundreds to thousands in a given metabolomics experiment, to find all detected drugs in a dataset.

Even when the drugs are identified, the interpretation of their biological impacts requires extensive literature and web searches to understand the therapeutic roles of the drugs and their mechanisms of action. Public databases, such as DrugBank,^16,17^ DrugCentral,^18^ DailyMed,^19^ and KEGG DRUG,^20^ can assist in interpretation, but the pharmacologic information is often provided as plain text or combinatorial classifications that require manual organization before downstream analysis. Although, in principle, large language models or similar text mining strategies can assist in this, the results of such models still need manual verification to confirm accuracy.^21–23^ In addition, it is not uncommon that only metabolized versions of a drug is present in the sample, leading to missed drug exposure if only the parent drug is considered.^24^ Unfortunately, with very few exceptions, reference MS/MS libraries include only the parent drug but not the drug metabolites due to challenges in obtaining reference standards for these metabolites.

The absence of MS/MS spectra for many drug metabolites, along with other challenges mentioned above, makes it very difficult to accurately annotate all drug exposures in biological specimens. For instance, stratifying a cohort based on antibiotic exposure - perhaps to better understand microbiome changes or as an exclusion criterion for clinical studies - requires identifying all antibiotics and their metabolites present in the samples. This is currently challenging due to the lack of resources that provide objective, systematic, and efficient readouts of drugs in untargeted metabolomics experiments.

To address this gap and to enable data science strategies on drug readouts, we curated the Global Natural Product Social Molecular Networking (GNPS) Drug Library, a collection of reference spectra for drugs and their metabolites/analogs (including parent ion masses and MS/MS spectra) along with structured pharmacologic metadata including exposure source, pharmacologic class, therapeutic indication, and mechanism of action. This comprehensive resource will enable further data science analysis to empirically - and retroactively - determine drug exposure using untargeted metabolomics data, complementing the information available in clinical records.

The creation of this library involved three key steps: 1) collecting MS/MS spectra of drugs and drug metabolites from publicly available MS/MS reference libraries; 2) finding MS/MS spectra analogs of those drugs in publicly accessible untargeted metabolomics data to enhance coverage of the metabolized versions of drugs; and 3) linking each MS/MS spectrum of a drug to controlled-vocabulary metadata - the key component of this resource that facilitates efficient data interpretation (**Figure 1a**).

**Figure 1.**
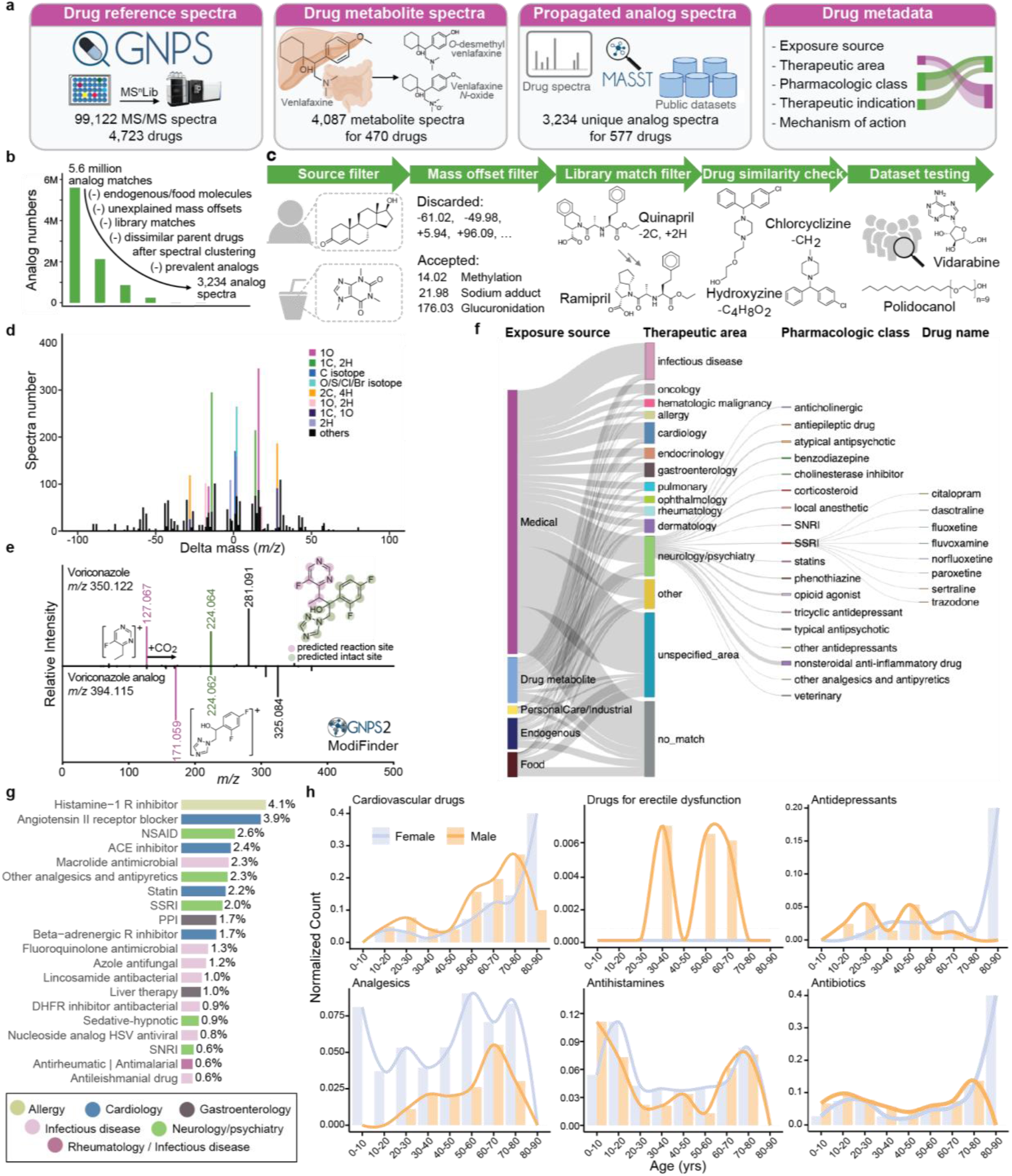
The GNPS Drug Library and connected pharmacologic metadata. **a**, The GNPS Drug Library comprises four key resources: Drug MS/MS reference spectra, drug metabolite MS/MS reference spectra, propagated drug analogs derived from public metabolomics datasets, and pharmacologic metadata connected to each reference spectrum. **b**, FastMASST analog search of drug spectra against public metabolomics studies yielded propagated drug-analogous MS/MS spectra, which were filtered by removing analogs for drugs with endogenous and food sources (source filter), removing mass offsets unexplained by common metabolic pathways (mass offset filter), removing analogs with GNPS library matches (library match filter), removing analogs connected to multiple drugs with dissimilar structures after spectra clustering (drug similarity filter), and removing analogs with unrealistic drug exposure indications (dataset testing). **c**, Illustration of each filter employed in curating FastMASST analog match results. **d**, Frequency of mass offsets in the propagated drug analog library. The mass offsets were grouped by unit mass and stacked based on the number of analog spectra. The most frequently observed mass offsets are colored while the rests are black. **e**, An example of structural modification sites predicted by ModiFinder.^32^ Purple color highlights modified spectra and substructures, while the green color highlights unmodified ones. **f**, Overview of the ontology-based drug metadata, highlighting common pharmaceutical classes and specific drugs in the neurology/psychiatry category. Width of the bars and lines reflects the number of unique drug structures in each class. **g**, The top 20 most detected pharmacologic classes in fecal samples from the American Gut Project,^33^ a cohort of the general population from the United States (US), Europe, and Australia (1,993 individuals). **h**, Detected therapeutic drug class patterns by age and sex (1,845 individuals with age and sex information; age 46 ± 18 years [range 3-93], with 53% being female). Detection of cardiovascular drugs increased with age, while analgesics, antihistamines, and antibiotics were detected across all ages.^34,35^ Analgesics were more frequently detected in females, consistent with the literature,^36,37^ and drugs for erectile dysfunction were detected only in males. NSAID, non-steroidal anti-inflammatory drugs; ACE, angiotensin converting enzyme; SSRI, selective serotonin reuptake inhibitor; PPI, proton pump inhibitor; DHFR, dihydrofolate reductase; HSV, herpes simplex virus; SNRI, serotonin and norepinephrine reuptake inhibitor.

The reference MS/MS spectra of drugs and their known metabolites were collected from two of the largest open-access mass spectral libraries, namely the GNPS Spectral Library^25^ and MS^n^Lib^26^. For all the MS/MS spectra in the GNPS and MS^n^Lib, metadata enrichment was first performed against PubChem (for synonyms),^27^ DrugCentral,^18^ the Broad Institute Drug Repurposing Hub databases,^28^ ChEMBL (for pharmacologic information),^29^ and DrugBank (for pharmacologic information and the Anatomical Therapeutic Chemical Classification code).^16,17,30^ This process utilized the available metadata in the GNPS Spectral Library and MS^n^Lib, including the chemical structures (e.g., SMILES or InChI), database identifiers (e.g., DrugBank ID or ChEMBL ID), and compound names. Based on the enriched metadata regarding clinical phases, all MS/MS spectra of drugs and compounds in clinical trials were compiled into the centralized GNPS Drug Library (see method details in **Supplementary Text 1**), resulting in 99,122 MS/MS reference spectra for 4,723 unique compounds. The compound names in the GNPS Drug Library were automatically curated and set to the first synonym in PubChem. We note that the term “drug” is used here in a broad sense, as the GNPS Drug Library includes not only prescribed and over-the-counter medications but also compounds currently in clinical trials, drugs that have been withdrawn, as well as substances with potential for abuse (e.g., cocaine, fentanyl).

Given that drug metabolites are largely overlooked in the initial search, we performed a second “partial name match” to include metabolites that retain the full drug names. For example, by searching for the name “venlafaxine” (a serotonin and norepinephrine reuptake inhibitor used to treat depression and various anxiety disorders), we obtained reference spectra for five of its metabolites, including “*N*-desmethylvenlafaxine”, “*O*-desmethylvenlafaxine”, “*N,O*-didesmethylvenlafaxine”, “*N,N*-didesmethylvenlafaxine”, and “venlafaxine *N*-oxide”. Using this strategy combined with manual inspection of the results, we captured 2,080 reference spectra for the metabolites of 110 drugs. Lastly, we added the MS/MS spectra collected in the development of dmCCS,^31^ a collision cross section database for drugs and their metabolites where human liver microsomes and S9 fraction were used for *in vitro* generation of drug metabolites. In total, 4,087 spectra for the metabolites of 470 drugs were included in the GNPS Drug Library (**Figure 1a**).

Despite the extensive collection effort, metabolite reference spectra were only available for 10% of the drugs included in the GNPS Drug Library. We hypothesized that unannotated drug metabolites are present in public untargeted metabolomics data. We further hypothesized that spectral alignment strategies can be used to find the modified versions of the drugs.^38–40^ In other words, public untargeted metabolomics data could be used to create a reference library of candidate drug metabolites that will facilitate the drug exposure readout in future datasets.

Based on MS/MS spectral alignment using two computational methods: repository-scale molecular networking^41^ and fast Mass Spectrometry Search Tool (fastMASST) with analog search,^42,43^ we retrieved all possible MS/MS spectra analogous to drugs from the GNPS/MassIVE public repository (∼2,700 LC-MS/MS datasets).^25^ These spectra represent drug-related molecules potentially derived from metabolism (host or microbiome), abiotic processes, and adducts of drugs from MS measurements. We obtained analogous MS/MS spectra for 14.6% of the 103,209 reference spectra for drug and drug metabolites (>5.5 million drug-analog spectral pairs).

In testing of the propagated analog library, we identified the need for additional filters to enhance its relevance to drug exposure (**Figure 1b-c**). First, it is not possible to determine the sources of exogenously supplied chemicals that are also produced endogenously or derived from the diet. Consequently, structural analogs of drugs with endogenous or dietary sources were excluded from the propagated drug analog library (e.g., analogous MS/MS spectra of testosterone used to treat hypogonadism, or caffeine used as a stimulant drug, were excluded). Second, propagated analogs with uncommon or unexplained mass offsets (precursor mass difference between the propagated analog and the connected drug) were excluded. Mass offsets were obtained from UNIMOD,^44^ from a community-curated list of explainable delta masses (**Table S1**), and from the Host Gut Microbiota Metabolism Xenobiotics Database,^45^ and were manually curated for those relevant to drug metabolism (e.g., 14.02 Da, methylation; 176.03 Da, glucuronidation) or mass spectrometry adducts (e.g., 17.03 Da, ammonium adduct; see **Table S2** for the 156 mass offsets that were included). MS/MS spectra of propagated analogs were excluded if the mass offsets were not in the customized list, or when the mass offsets occurred fewer than ten times. Third, since drugs within the same pharmacologic family often have similar structures, they can be identified as analogs of each other through spectral alignments. Therefore, we excluded MS/MS spectra with matches to the GNPS library from the propagated analog annotations. For example, a propagated analog of quinapril, an angiotensin converting enzyme (ACE) inhibitor, had a spectral match to ramipril, another ACE inhibitor (**Figure 1c**). Excluding these analog annotations ensures that they do not overwrite library matches of known drugs and metabolites. Fourth, if one propagated analog spectrum is connected to multiple drugs after spectral clustering, the drugs need to be structurally similar to accept the shared analog. We illustrate this with the propagated analog (*m/z* 287.133, with a formula C_17_H_19_ClN_2_) that is connected to both hydroxyzine [C_21_H_27_ClN_2_O_2_, mass offset 88.05 Da (C_4_H_8_O_2_)] and chlorcyclizine [C_18_H_21_ClN_2_, mass offset 14.02 Da (CH_2_)], which share the core structure (**Figure 1c**). Finally, we tested the propagated drug analog library against 12 public LC-MS/MS datasets to filter out analogs that have unrealistic drug exposure indications. The selected datasets represent a broad range of human tissue types and biofluids, including fecal (n=5), breast milk (n=2), plasma (n=3), skin (n = 1), and brain (n=1), as well as multiple mouse tissues (n = 2; with metadata confirming no drugs were used). We observed analogs of tocofersolan (a synthetic vitamin E derivative), iloprost (a synthetic prostacyclin mimetic), desonide (a synthetic topical corticosteroid), medroxyprogesterone (a synthetic progestin), and vidarabine (an adenosine analog used as an antiviral) in >50% of the human fecal samples from the American Gut Project (n = 1,993 individuals), a cohort of the general population. The connected drugs for these analogs are derivatives of endogenous or food derived molecules and are unlikely to be used by more than half of the population. Therefore, these analogs cannot be confidently linked to drug exposures and were excluded. Analogs of polidocanol (a synthetic long-chain fatty alcohol used as anesthetics) were observed in >70% of 2,463 human milk samples. They are likely surfactants/contaminants with the polyethylene glycol structural units^46^ and thus were excluded from the propagated drug analog library (**Figure 1c**).

After all filtering steps, 3,234 clustered MS/MS spectra representing propagated analogs of 577 drugs were retained in the final drug analog library. We observed that 75% of the propagated analogs occurred at least once in the same data file with the corresponding parent drugs (**Table S3**). The most common mass offsets in the drug analog library correspond to a gain of 15.99 Da, which can be interpreted as the gain of an oxygen (e.g., oxidative metabolism of the drug), followed by a gain or loss of 14.02 Da (CH_2_, (de)methylation), a gain of 1.00 Da (13C isotope), a gain of 2.00 Da (O/S/Cl/Br/2C isotopes), and a gain or loss of 28.03 Da (C_2_H_4_, (de)ethylation; **Figure 1d**). Notably, it is possible that such analog spectra are MS/MS of other ion forms of the parent drug, such as isotopes, different adducts, in/post-source fragments, or multimers, rather than drug metabolites or structural analogs. However, their indications in drug exposure remain the same and thus we did not separate drug metabolites and instrument adducts in drug exposure stratification. To extend structural hypotheses for the drug analogs, we employed the newly developed ModiFinder,^32^ which leverages the shifted MS/MS fragment peaks in the MS/MS alignment to predict the most likely location of the structural modifications. We were able to predict the partial location of the modification for 61.5% of the analog spectra. We demonstrate examples where ModiFinder predictions agree with expert manual interpretation of the MS/MS spectra (**Figure 1e, S1**).

Connecting drug detections to their therapeutic indications typically requires expert knowledge and/or extensive literature searches. The GNPS Drug Library addresses this challenge by providing controlled-vocabulary metadata together with the specific drug annotations. This allows users to annotate all drugs in an untargeted metabolomics dataset and directly obtain a table with exposure sources, pharmacologic classes, therapeutic indications, and mechanisms of action of the drugs, with their structures and names in a data science ready format (**Figure 1f, S2**). Particularly, the “exposure source” information categorizes the drugs in a combination of five classes, namely medical, endogenous, food, personal care, and industrial sources, which was based on the source categorizations from the Chemical Functional Ontology (ChemFOnt) database^47^ and modified manually - by parsing of web pages and scientific literature - to increase compound coverage and improve accuracy and consistency. This categorization allows distinguishing endogenous or food sourced molecules (for the non-analogous spectra only). Examples include deoxycholic acid, an endogenous molecule also used for liver disease, and lactitol, a food sweetener also used as a laxative. Using the GNPS Drug Library metadata, such annotations can be separated from those molecules used exclusively as drugs, which have entirely different exposure implications.

Through structural and name matches, we extracted the pharmacologic classes of 900 drugs from the U.S. Food and Drug Administration (FDA) and the therapeutic areas, therapeutic indications, and mechanisms of action for 3,894 drugs from the Broad Institute Drug Repurposing Hub.^28^ However, we noticed substantial variability in the extracted information (e.g., inconsistent therapeutic areas assigned to drugs within the same pharmacological class; the sulfonamide antimicrobials sulfamethizole, sulfamethazine, and sulfacetamide were categorized as infectious disease, gastroenterology, and ophthalmology, respectively), or insufficient metadata for several drugs (e.g., common therapeutic indications missing). Therefore, this metadata was further manually curated by expert clinical pharmacologists to enhance and clean up the information retrieved from databases. This manual curation increased the metadata coverage to 4,560 drugs. Drugs without associated metadata are typically those that have been withdrawn from the market (e.g., indoprofen), were in drug development but never marketed (e.g., tarafenacin), or are under development but do not yet have regulatory approval (e.g., firsocostat).

In total, 735 drugs in the GNPS Drug Library (38,001 spectra) were identified with endogenous or dietary sources. The final metadata of the drug library covers 27 unique therapeutic areas, 571 pharmacological classes, 920 therapeutic indications, and 823 mechanisms of action (**Figure 1f, S2**). Therapeutic areas of neurology/psychiatry, infectious disease, and cardiology have the highest number of included drugs (**Figure 1f**) and reference spectra (**Figure S2**). We note that these incidences reflect the availability of the reference spectra but not the prevalence of these drugs in the general population. Combining the exposure source and therapeutic area, we noticed that fewer drugs related to infection and neurology/psychiatry have endogenous or food sources, while higher portions of drugs used for gastroenterology (e.g., deoxycholic acid, riboflavin) and dermatology (e.g., salicylic acid, nicotinamide) are endogenous and/or can come from food-derived molecules.

In order to assess the utility of the GNPS Drug Library for detecting drugs known to be consumed, we analyzed two pharmacokinetic datasets where healthy individuals received certain drugs followed by time-series sampling. In the first study, 10 participants received a single oral dose of diphenhydramine.^48^ The drug was not detected in plasma and skin samples before administration, but was detected in all individuals post-administration over the course of 24 hours (**Figure S3a**). In plasma, detection frequencies peaked at 1-2 hours (**Figure S3a**), aligning with the reported time to maximum concentration (∼2 hours) for diphenhydramine.^49^ In skin, peak detection occurred at 10-12 hours (**Figure S3a**), reflecting the delayed deposition to skin compared to plasma for orally administered drugs. In the second study, 14 participants received a cocktail of oral caffeine, midazolam, and omeprazole.^50^ These drugs were detected in plasma from 100%, 69%, and 100% of participants, respectively (**Figure S3b**). The detection frequencies in fecal samples were below 25%. The same participants began a 7-day course of oral cefprozil (administered twice daily; day 2-8) the day after the cocktail drug administration. Cefprozil was detected in fecal samples with increasing frequencies from 0% at day 2 to 43% at day 9. These results demonstrate that the GNPS Drug Library can reliably detect consumed drugs and that detection is both biofluid and time dependent. The results also emphasize the need to establish medication exposures empirically in the context of the analyzed samples, as clinical studies rarely recorded the time between drug intake and sample collection.

Connected with public untargeted metabolomics data, the GNPS Drug Library can reveal distinct drug exposure profiles among individuals with different disease, age, and sex. For different disease studies, we used the human disease ontology identifier (DOID) curated in ReDU, a controlled-vocabulary metadata for public metabolomics datasets,^51^ and searched for the drugs and drug analogs using fastMASST.^42^ Samples from individuals with inflammatory bowel disease, Kawasaki disease, and dental caries were characterized by high detection frequencies of antibiotics (**Figure S4a**). Skin swabs of patients with psoriasis were characterized by antifungals. Samples from people with human immunodeficiency virus (HIV) showed high frequency of antivirals, and samples from individuals with Alzheimer’s disease were characterized by cardiology and neurology/psychiatry drugs, all consistent with the expected drug usage of people with these diseases (**Figure S4a**).

To investigate drug exposures among different age and sex groups, we profiled 1,993 fecal samples from the American Gut Project,^33^ with participants from the United States (US), Europe, and Australia with age 46 ± 18 years (range 3-93; 53% female). A total of 75 different drugs were detected; the most frequently detected pharmacologic classes included histamine-1 receptor antagonist (allergy), angiotensin II-receptor blocker (cardiology), ACE inhibitor (cardiology), beta-adrenergic receptor inhibitor (cardiology), statin (lipid-lowering), non-steroidal anti-inflammatory drug (NSAID; analgesics), and selective serotonin reuptake inhibitor (SSRI; antidepressant), which matches with the most commonly prescribed drug classes in these regions (**Figure 1g**).^52–54^ There were more drugs per individual noted in the US cohort compared to the European and Australian cohorts (chi-square test; χ^2^ (8, n = 1,903) = 33, p = 5.3 × 10^−5^, **Figure S4b**). When connected with age and sex information, the drug detection agrees with the expected usage patterns of different drug classes (**Figure 1h**). For example, cardiovascular drugs were detected more frequently with increasing age, while analgesics, antihistamines, and antibiotics were detected across all ages.^34,35^ We also observed that analgesics, such as NSAIDs and paracetamol, were more frequently detected in females (chi-square test; χ^2^ (1, n = 1,958) = 15.4, p = 8.54 × 10^−5^), consistent with the literature,^36,37^ and that drugs for erectile dysfunction were detected only in males. Overall, empirical drug readout using untargeted metabolomics, facilitated by the GNPS Drug Library, demonstrated good specificity among individuals with different disease, age, and sex.

The GNPS Drug Library can allow the discovery of previously uncharacterized drug metabolites. To illustrate this, we analyzed fecal samples from the HIV Neurobehavioral Research Center (HNRC) cohort (n = 322; age 55 ± 12 years), which included both people with HIV (n = 222) and people without HIV (n = 100). Among the 17,729 unique MS/MS spectra obtained, 493 were annotated with the GNPS Drug Library. After removing drugs that could be from endogenous or food sources (because we cannot assess whether they were given as a medication) and grouping annotations of drugs, metabolites, or analogs, 169 unique drugs remained. Antiretroviral drugs (ARVs; drugs for the treatment of HIV), drugs for cardiovascular disease, and drugs for anxiety and depression were the most frequently detected categories (**Figure 2a, S5a**). Despite the high rates of viral suppression with the advent of antiretroviral therapies (ART; a combination of ARVs to treat HIV), people with HIV have disproportionately high rates of depression and cardiovascular diseases,^55–58^ reflected in the observation of antidepressants and cardiovascular drugs in these samples.

**Figure 2.**
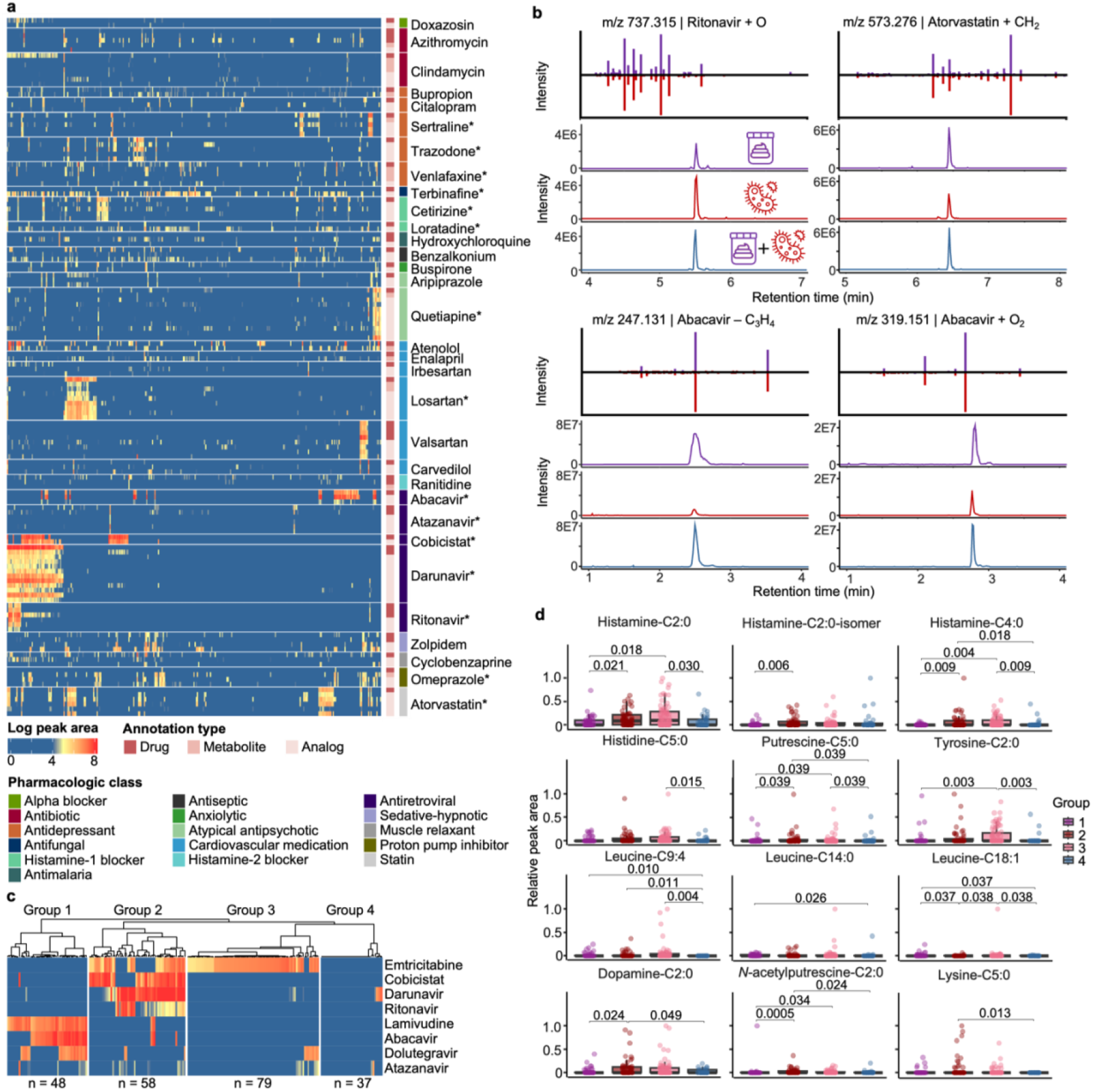
Drug exposures in the HIV Neurobehavioral Research Center (HNRC) cohort with connections to microbial metabolism and endogenous metabolites. From the HNRC cohort, 322 fecal samples were analyzed with 222 samples from people with HIV and 100 samples from people without HIV. **a**, Peak area visualization of drugs detected with metabolites and analogs. Each column represents one sample and each row represents one drug annotation. Drug annotations were grouped based on the parent drugs and separated by gap spaces. Drug annotations were denoted based on their types (as drug, drug metabolites, or drug analogs) and the pharmacologic classes of the parent drugs. All annotated ion/adduct forms of the parent drugs were visualized, leading to multiple rows of parent annotations for some drugs. Asterisks on the drug name mark parent drug annotations confirmed with commercial standards based on retention time and MS/MS spectral matches. Raw peak areas were log-transformed. **b**, Retention time and MS/MS spectra mirror matches for drug analogs observed in both the fecal samples and the drug microbial incubations. Purple traces represent the fecal samples, while red traces represent the drug microbial incubation. Blue traces represent mixtures of the fecal samples and the microbial incubations at 1:1 volume ratio. The atomic changes of the drug analogs were based on [M+H]^+^ ion of the parent drug. **c**, Hierarchical clustering of the samples from people with HIV (n = 222) based on detected antiretroviral drugs (ARV). Each row represents one detected ARV, with peak areas summed for the drug, metabolite, and analog detections followed by log-transformation (visualized with the same color scale as panel a). ARVs detected in <10% of samples are not shown. Each column represents one sample, clustered into four groups by hierarchical clustering with Ward’s linkage and Euclidean distance. **d**, Sample-to-sample peak areas of *N*-acyl lipids in people with HIV, separated by the clusters derived from ARV detections shown in panel c. For each compound, the peak area in each sample was standardized to the maximum value observed across all samples. A non-parametric Kruskal-Wallis test followed by pairwise Wilcoxon test and Benjamini-Hochberg correction for multiple comparisons were performed. P-values < 0.05 were noted in the figure. Boxplots showcase the median value, first (lower) and third (upper) quartiles, and whiskers indicate the error range as 1.5 times the interquartile range.

Interestingly, 33% of the drugs were annotated together with their metabolites or analogs, and the occurrences of drug metabolites/analogs aligned with those of the parent drugs (**Figure 2a**). For example, darunavir (an ARV) had no annotated metabolites but was observed with 10 analogs (**Figure S5b**). Retention time and peak shape analysis indicated that two of the darunavir analogs are in-source fragments (as judged by overlapping retention times),^59^ while the others remain unknown metabolites or isomers of this drug (**Figure S5c**,**d**). For the analogs that are not in-source fragments, 63-100% (median 96%) of their occurrences were together with the darunavir parent drug. The observations of darunavir analogs without the parent drug are perhaps related to the timepoint of sample collection or to individuals with an ultrarapid metabolizer phenotype, impairing the detection of the parent drug. Nevertheless, this observation highlights the utility of drug metabolites and analogs to increase the sensitivity of drug exposure readouts via untargeted metabolomics. We note that the HNRC dataset was added to the GNPS/MassIVE public repository after the development of the drug analog library. Therefore, analog mining via existing public metabolomics datasets can facilitate the discovery of uncharacterized metabolites in new data.

To further investigate the potential metabolic sources of the observed drug analogs, we cultured darunavir and 12 other drugs with a defined and complex synthetic microbial community of 111 bacterial species commonly found in the human gut.^60^ Except clindamycin (an antibiotic), all drugs observed with three or more metabolites/analogs that were present in >10% samples were incubated (10 drugs in total; **Table S4**); omeprazole, loratadine, and terbinafine were additionally included because their analogs were frequently observed in samples without the respective parent drugs. Shared analogs were observed for 10 of the 13 drugs between the fecal samples and the microbial incubations. Among them, metabolites/transformation products were observed for 4 drugs (ritonavir, atorvastatin, abacavir, and omeprazole; **Figure 2b, S6**), while the rest of the analogs were in-source fragments based on retention time correlation analysis.^59^ The ritonavir, atorvastatin, and abacavir analogs increased in intensity with increased microbial incubation time (**Figure S6a-d**), indicating microbial metabolism as a possible source and consistent with their observation in fecal samples. The omeprazole analog (*m/z* 330.127) appeared to be an abiotic transformation product because it was already present at t=0 cultures, and its intensity decreased with increased incubation time (**Figure S6e-g**). This is consistent with the fast activation of omeprazole (*m/z* 346.122), a proton-pump inhibitor and a prodrug, to the reactive sulphenamide product (*m/z* 330.127) at low pH.^61^ Rapid photolysis and hydrolysis of omeprazole has also been reported in abiotic environments with a major deoxygenation transformation product (*m/z* 330.127).^62,63^

The GNPS Drug Library can enable stratification based on drug profiles, which facilitates discovery of connections between drug exposures and endogenous metabolites. *N*-acyl lipids are a class of signaling molecules made by host-associated microbiota^64^ that play important roles in the immune system,^65^ memory function,^66^ and insulin regulation of the human body.^67–69^ Our recent ongoing work found that the levels of histamine *N*-acyl lipids differed by HIV serostatus. Specifically, we observed higher levels of histamine-C2:0, histamine-C3:0, and histamine-C6:0 in people living with HIV than people without HIV.^70^ To investigate whether these differences were related to drug exposures, we further stratified samples in this dataset by their ARV exposure profiles. As expected, the ARV profiles clearly separated based on the HIV serostatus (**Figure S7a**). High intensities of different ARVs were observed in fecal samples from people with HIV, while ARVs were only occasionally observed in people without HIV with low intensities. ARVs observed in people without HIV include tenofovir, maraviroc, atazanavir, and raltegravir, which are commonly used for prophylaxis (**Figure S7a**).^71,72^ To control for the HIV serostatus and investigate the effects of ARV exposure, we excluded samples from people without HIV and stratified the people with HIV (n = 222) based on their ARV co-occurrences. Four distinct ARV exposure groups were observed based on hierarchical clustering that agreed well with the different combination antiretroviral therapy (cART) regimens (**Figure 2c**). For example, Group 1 (n = 48), characterized by lamivudine, abacavir, and dolutegravir exposures, corresponded to the dolutegravir/abacavir/lamivudine treatment regimen.^73^ Group 2 (n = 58) with emtricitabine, darunavir, ritonavir, and cobicistat exposures, agreed with the darunavir/ritonavir regimen^74^ and the darunavir/cobicistat/emtricitabine/tenofovir regimen.^75^ Group 3 (n = 79), characterized by emtricitabine and dolutegravir exposures, may be related to the dolutegravir/emtricitabine/tenofovir treatment regimen (Group 2).^76^ Group 4 (n = 37) were without apparent ARV exposures, possibly due to poor adherence, severe comorbidities, HIV elite control, or ARVs not included in the GNPS Drug Library or not amenable with LC-MS/MS detections (**Figure 2c**). Notably, we observed that the levels of histamine-C2:0 previously associated with HIV serostatus,^70^ along with the levels of eleven other *N*-acyl lipids, were significantly different in the four ARV exposure groups (Kruskal-Wallis test, p-value < 0.05; see specific p-value in **Figure 2d**). This suggests that exposure to different classes of ARV among people with HIV, in addition to HIV serostatus itself, might contribute to changed levels of these *N*-acyl lipids. We emphasize that these patterns could not have been revealed without the empirical drug readouts from untargeted metabolomics. Clinical research records may not document exposures to individual drugs and often do not provide quantitative information on the exposure levels. For example, metadata for the HNRC cohort on current ARV usage, which is based on self-reports, documented drug usage as “ARV-naïve” (never received ARV), “no ARV” (no current ARV use), “non-HAART” (currently using less than three ARVs), and “HAART” (currently using three or more ARVs). Based on these classifications, no significant differences were observed for the 52 *N*-acyl lipids detected in these samples (**Figure S7b**). Without the empirical drug readout, enabled by the GNPS Drug Library, the effects of drugs on microbial *N*-acyl lipid levels would be overlooked.

We anticipate the GNPS Drug Library to play a key role in precision medicine by enhancing our understanding of the effects of drugs across a wide range of phenotypes, including endogenous metabolism, gut and skin microbiome, pharmacokinetics, and drug-drug interactions. The empirical drug readouts from the GNPS Drug Library can enhance the clinical metadata by providing sample-to-sample comparisons of the relative abundance of individual drugs, which can be flexibly summarized at multiple ontology levels depending on user-defined questions. The mass spectrometry community will play a key role in the evolution of this resource through the continued deposition of reference libraries and expansion of the public metabolomics datasets for analog searches. By harnessing the power of public data and data science-ready metadata, we can unlock opportunities to deepen our understanding of the intricate relationships between xenobiotic exposure and human biological systems.

It is important to understand that the use of the GNPS Drug Library holds certain limitations. The current library only supports MS/MS-based annotations to level 2/3 according to the 2007 Metabolomics Standards Initiative.^77^ This generally means that spectra of drug isomers may be annotated as the drug. Key drugs with important clinical implications should be checked for retention time matching and be quantified with analytical standards should the scientific question warrant this. The GNPS Drug Library can only capture drugs that are detectable in the specific biological matrix of choice (e.g., brain samples and urine will likely have different drug exposure readouts) and drugs that are ionizable with the chosen mass spectrometry setup. When constructing the drug analog library, we designed the filters to retain analog spectra that can be as confidently linked to drug exposure as possible, at the likely cost of excluding true positives. For example, metabolism pathways specific to substructures infrequently captured or missing in our customized delta mass list will be excluded. Metabolites shared between drugs that cannot be connected by the applied structural similarity scores (the Tanimoto score)^78^ will be lost. As this is an evolving resource, we encourage the community to not only add to, but also report any inconsistencies in the library and the metadata they may notice.

## Acknowledgements

This project was enabled in part by the Alzheimer’s Gut Microbiome Project (AGMP) and the Data Infrastructure and Molecular Atlas for AD: Connection Exposome, Gut Microbiome, and Metabolome supplement funded wholly or in part by the following grants thereto: 1U19AG063744 and 3U19AG063744-04S1 and awarded to Dr. Kaddurah-Daouk at Duke University in partnership with multiple academic institutions. As such, the investigators within the AGMP and the Exposome Supplement, not listed specifically in this publication’s author’s list, provided data along with its pre-processing and prepared it for analysis, but did not participate in analysis or writing of this manuscript. A listing of AGMP Investigators can be found at https://alzheimergut.org/meet-the-team/. A complete listing of ADMC investigators can be found at: https://sites.duke.edu/adnimetab/team/. We also thank the support by NIH for the Maternal and Pediatric Precision in Therapeutics project P50HD106463, the development of tools for structure elucidation R01DK136117, the Collaborative Microbial Metabolite Center U24DK133658. The HIV Neurobehavioral Research Center (HNRC) is supported by Center award P30MH062512 from NIMH. This research was supported in part by the Intramural Research Program of the NIH, National Institute of Environmental Health Sciences (ZIC ES103363). C.B. was supported by the Czech Academy of Sciences PPLZ fellowship number L200552251. V.C.L is supported by Fonds de recherche du Québec - Santé (FRQS) Postdoctoral fellowship (335368). N.E.A was supported in part by the National Center for Complementary and Integrative Health of the NIH under award number F32AT011475. A.M.C.-R. and P.C.D. were supported by the Gordon and Betty Moore Foundation grant GBMF12120. M.R. was supported by the NIH grant R37 AI126277. T.P. was supported by the Czech Science Foundation (GA CR) grant 21-11563M and by the European Union’s Horizon 2020 research and innovation programme under Marie Skłodowska-Curie grant agreement No. 891397.

## Author contributions

H.N.Z., C.B., P.C.D. conceptualized the method. H.N.Z, C.B., R.S., T.P. developed the MS/MS library for drugs. H.N.Z., W.B., R.T., R.S. developed the MS/MS library for drug analogs. H.N.Z., S.M., H.S., P.R., curated exposure source metadata. K.E.K. curated pharmacological metadata. H.N.Z., R.T., Y.E.A., H.M.-R., N.E.A., A.M.C.-R., P.W.P.G., S.M., I.M., A.C., S.P.T., S.Z., curated drug name match results. H.N.Z., K.E.K., S.L., H.M.-R., L.K., S.X., performed data analyses. Y.E.A., M.S.A., C.W., A.K.J., D.M., helped with data interpretation. V.C.-L., H.N.Z., L.C., C.W., M.R., K.Z. performed microbial incubation experiments. D.H.R., L.X. contributed MS/MS reference spectra. M.S.A., R.J.E., D.F., R.H., J.I., S.L., D.J.M. developed the clinical cohort of human immunodeficiency virus (HIV) infection. M.R.Z.S., M.W. performed ModiFinder analysis. S.G., D.S.W. provided support on exposure source annotation. R.K.-D. supervised the consortium providing access to the Alzheimer’s disease cohort and acquired funding. R.K. supervised sample handling and DNA data acquisition for the Alzheimer’s disease, HIV and American Gut Project cohorts, and acquired funding. H.N.Z., K.E.K., C.B., P.C.D. drafted the manuscript. P.C.D., S.M.T. acquired funding and supervised this project. All authors reviewed and edited the manuscript.

## Data availability

The MGF spectral files for the GNPS Drug Library and the associated metadata of controlled vocabularies (.csv) can be downloaded from Zenodo archive under doi: 10.5281/zenodo.13892289. The downloaded MGF spectral files can be added to personal GNPS folders and used directly for library matching. Data used to validate the empirical drug readouts are publicly available in GNPS/MassIVE under the accession numbers MSV000085944, MSV000084008, and MSV000082493. Data used to profile drug exposures by age and sex are available at MSV000080673. Data for fecal samples from the HNRC cohort are available at MSV000092833. Data for the drug bacterial cultures are available at MSV000095331. Data for HNRC fecal samples analyzed with the bacterial cultures are available at MSV000096012. Data for co-migration of the bacterial cultures and fecal samples are available at MSV000096013. Due to human subject protection constraint, metadata for the HNRC cohort will be provided upon request to HNRC: https://hnrp.hivresearch.ucsd.edu.

## Code availability

The code used to query reference spectra of drugs is available on GitHub under the MS^n^ library project^26^ (https://github.com/corinnabrungs/msn_tree_library). The code used to filter the drug analog matches is provided on GitHub (https://github.com/ninahaoqizhao/Manuscript_GNPS_Drug_Library). The code used for dataset analysis can be found on GitHub (https://github.com/ninahaoqizhao/Manuscript_GNPS_Drug_Library and https://github.com/kinekvitne/manuscript_drug_library).

## Supplementary Figure

Supplementary Figure S1. Additional examples of structural modification sites of the drug analogs predicted by ModiFinder; Supplementary Figure S2. Overview of the ontology-based drug metadata based on the numbers of reference spectra; Supplementary Figure S3. Empirical drug readout in healthy individuals receiving specific drugs; Supplementary Figure S4. Drug exposure profiles among cohorts with different diseases and geolocations by re-analyzing public metabolomics data; Supplementary Figure S5. Drug analog annotations in fecal samples from people with human immunodeficiency virus; Supplementary Figure S6. Drug analogs observed in human fecal samples can be produced by microbial metabolism; Supplementary Figure S7. Comparison of sample clustering based on empirical drug records from the GNPS Drug Library or on clinical metadata; Supplementary Figure S8. Retention time and MS/MS spectra mirror matches for drugs observed in the HNRC cohort with analytical standards.

## Supplementary Table

Supplementary Table S1. Community-curated list of delta mass interpretation; Supplementary Table S2. Delta masses accepted in the drug analog library; Supplementary Table S3. Percentage of drug analogs demonstrating co-occurrence with the parent drugs in MASST search, separated by delta masses; Supplementary Table S4. Source of drugs used in synthetic microbial community incubation; Supplementary Table S5. Bacterial strains used in the six synthetic microbial communities; Supplementary Table S6. Composition of the BHI medium for anaerobic microbial cultures.

