## Supplementary Methods for "Empirically establishing drug exposure records directly from untargeted metabolomics data"

[19] Department of Medicinal Chemistry, University of Washington, Seattle, WA, USA
[20] Current address: Biological Sciences Division, Pacific Northwest National Laboratory, Richland, WA, USA
[21] Department of Computer Science and Engineering, University of California Riverside, Riverside, CA, USA
[22] Center for Microbiome Innovation, University of California San Diego, La Jolla, CA, USA.
[23] Department of Computing Science, University of Alberta, Edmonton, AB T6G 2E8, Canada
[24] Department of Psychiatry and Behavioral Sciences, Duke University, Durham, NC, 27708, USA
[25] Duke Institute of Brain Sciences, Duke University, Durham, NC, USA
[26] Department of Medicine, Duke University, Durham, NC, USA
[27] Chiba University, UC San Diego Center for Mucosal Immunology, Allergy, and Vaccines (CU-UCSD cMAV), La Jolla, CA, USA
[28] Program in Materials Science and Engineering, University of California, San Diego, 9500 Gilman Drive, La Jolla, CA 92093-0418, USA
[29] Department of Computer Science and Engineering, University of California San Diego, La Jolla, CA, USA
[30] Shu Chien-Gene Lay Department of Bioengineering, University of California San Diego, La Jolla, CA, USA
[31] Halıcıoğlu Data Science Institute, University of California San Diego, La Jolla, CA, USA
[†] Haoqi Nina Zhao, Kine Eide Kvitne, and Corinna Brungs contributed equally to this work.
* Author to whom correspondence should be addressed.

### ***Reference spectra collection and curation***

Reference spectra of drugs were collected from the GNPS spectral library[25] and the $MS^n$ library using a Python script developed for the $MS^n$Lib workflow.[26] Detailed steps are summarized in the **Supplementary Text 1**. Briefly, the chemical structures provided in the open-source libraries are first cleaned by removing salt forms and standardized with the ChEMBL structure pipeline Python package.[79] Based on this cleaned structure, other structural information is calculated, including the canonical and, if available, isomeric SMILES, InChI, and InChIKey strings. Database identifiers, e.g., DrugBank ID or ChEMBL ID, are searched in PubChem and Unichem based on the InChIKey string. Those identifiers, the complete InChIKey or the first part of the InChIKey to remove stereochemistry, are then used to search in drug databases, including the Broad Institute Drug Repurposing Hub (downloaded July 2022) ,[28] DrugBank (downloaded August 2022),[16,17] and DrugCentral (downloaded October 2022),[18] and the database of bioactive molecules from ChEMBL with clinical phases 1-4 (downloaded March 2023).[29] Reference spectra with chemical structures contained in any of the databases above are retained in the GNPS Drug Library.

To increase the coverage for drug metabolites and derivatives, we searched for reference spectra whose name contained the full name of a drug in the abovementioned databases, followed by manual inspection to remove mismatches. For example, by searching for the name "venlafaxine" (a serotonin and norepinephrine reuptake inhibitor indicated for the treatment of depression), we collected reference spectra for five metabolites of venlafaxine, including "*N*-desmethylvenlafaxine", "*O*-desmethylvenlafaxine", "*N*,*O*-didesmethylvenlafaxine", "*N*,*N*-didesmethylvenlafaxine", and "venlafaxine *N*-oxide" from the GNPS Library.

### ***Drug analog search and result filtering***

Drug analog spectra were retrieved from the GNPS/MassIVE public repository based on MS/MS spectra alignment using two computational methods: repository-scale molecular networking[41] and fastMASST with analog search.[42,43] A repository-scale molecular network was constructed in our previous research[41] and annotated with the 103,209 reference spectra of drugs and drug metabolites collected in this study. Unannotated spectra directly linked to annotated drugs or drug metabolites were collected as tentative drug analogs. FastMASST (fast Mass Spectrometry Search Tool)[42,43] checked the similarity of the queried spectrum against all spectra in the public repository without the pre-constructed molecular network. Batch mode FastMASST search for the 103,209 reference spectra of drugs and drug metabolites was achieved in July 2023 with the customized Python scripts (https://github.com/robinschmid/microbe_masst), leveraging the Fast Search tool API.[80] The Python scripts returned analog matches as metabolomics universal spectrum identifiers (USIs),[81] connecting to spectra in the GNPS/MassIVE repository. Analog spectrum matches were accepted with a modified cosine score of 0.8 or higher and a minimum of 6 matching ions, with a search window of 150 Da below and 200 Da above the precursor mass.

Filtering of the analog matches was achieved using customized Python scripts with the following steps. *Step One,* analog matches for drugs with endogenous or food sources, based on the manually curated metadata, were excluded to avoid matches to endogenous metabolites. *Step Two,* analogs with mass offsets (precursor mass differences between the analogs and the drugs) unrepresented by common metabolism pathways were excluded. Specifically, mass offsets sourced from UNIMOD,[44] from a community-curated list of delta masses (**Table S2**), and from the Host Gut Microbiota Metabolism Xenobiotics Databases[45] were manually curated to only keep those relevant to drug metabolism (e.g., 14.02 Da, methylation; 176.03, glucuronidation) or mass spectrometry adducts (e.g., 17.03, ammonium adduct; see **Table S3** for the 156 mass offsets included). Analogs were excluded if the mass offsets were not in the customized list, or if the mass offsets occurred fewer than ten times. *Step Three,* analogs with matches to the GNPS Library were excluded to restrict the analog library to true unknowns. This is achieved through fastMASST searches of the GNPS Library (analog search off, minimum cosine score 0.7, minimum 5 matching ions); all USIs with library matches were excluded from the drug analogs. *Step Four,* analogs matched to multiple drugs with dissimilar structures were excluded. Specifically, spectra of the remaining USIs from Step Three were downloaded using the metabolomics-usi API[81] and clustered by the falcon spectrum clustering tool (with default setting except an eps score of 0.05).[82] Singleton spectra were excluded. For clustered spectra connected to multiple drugs, Tanimoto similarities of the drugs need to be higher than 0.8 to accept the drug analogs.[78] *Step Five,* the drug analog libraries were tested against 10 datasets to remove edge cases where the analogs were detected with high frequency and cannot be confidently linked to drug exposure. The testing datasets include four datasets for human fecal samples (GNPS/MassIVE ID: MSV000080673, MSV000095418, MSV000094515, MSV000092833), two for human breast milk samples (MSV000091520, MSV000090877), one for human plasma (MSV000094395), one for human brain tissues (MSV000086415), and two for mouse tissues (MSV000091007, MSV000091363). Search against the GNPS Drug Library was performed using the Library Search workflow on GNPS with minimum cosine score of 0.7 and minimum 5 matching peaks.

To extend structural hypotheses for the drug analogs, we employed the newly developed ModiFinder,[32] which leverages the shifted MS/MS fragment peaks in the MS/MS alignment to predict the most likely location for the structural modifications. ModiFinder was run with mass tolerance of 40 ppm and fragmentation depth of 2. The GNPS job is available at: https://gnps2.org/result?task=f4bf3bab255c4bd8a4175433286d5f2c&viewname=modifinder_result&resultdisplay_type=task.

***Metadata curation***

The GNPS Drug Library provides ontology-based metadata on the chemical sources, therapeutic areas, pharmacologic classes, therapeutic indications, and mechanisms of action of the drugs. The chemical source information was first retrieved from the ChemFOnt database[47] through name matching. Synonyms of the drugs in the GNPS Drug Library were extracted from PubChem with the $MS^n$ library Python scripts and searched against chemical names in the ChemFOnt database. The results were manually curated by three experts through parsing web pages and scientific literature to increase compound coverage (all drugs in the GNPS Drug Library have chemical source information) and improve accuracy and consistency. The therapeutic areas, mechanisms of action, and therapeutic indications of the drugs were extracted from the Broad Institute Drug Repurposing Hub[28] using the $MS^n$ library Python scripts through InChIKey matches. The pharmacologic classes were extracted from the U.S. Food and Drug Administration (FDA) documents by synonym matching. The above database search returns the therapeutic areas, mechanisms of action, and therapeutic indications of 3,894 drugs (86,364 spectra) and the pharmacologic classes of 900 drugs (22,935 spectra). The extracted metadata were manually curated by expert pharmacologists to improve accuracy and fill in missing information to the extent possible, which increased the metadata coverage to 4,560 drugs (90,325 spectra). Sources used to curate the metadata included the Anatomical Therapeutic Chemical (ATC) Classification, National Institutes of Health (NIH), FDA, European Medicines of Agency (EMA), and DrugBank. In Particular, the WHO ATC codes were used in the manual curation of the pharmacologic class. The refined metadata can be illustrated with doxycycline, a tetracycline-class drug. Initially, it was classified under dental as a therapeutic area with periodontitis as the only indication. Following manual curation, doxycycline was reclassified under infectious diseases (therapeutic area) and Gram-positive and Gram-negative infections (therapeutic indication), recognizing its usage to treat a variety of infections. For drug metabolites and analogs, metadata corresponding to the parent drug was used.

***Ethics oversight***

All datasets included in the manuscript were approved by the Institutional Review Board for Human Research, either by the University of California San Diego or University of Colorado, Boulder, and performed in accordance with the Declaration of Helsinki: American Gut Project, protocol no. 141853 and 12-0582; diphenhydramine study, protocol no. 191026; Cooperstown cocktail study, protocol no. 161940; HNRC, 172092. All subjects provided written informed consent.

***Reanalysis of data from pharmacokinetic studies with known drug administration***

In order to validate the empirical drug records from GNPS Drug Library, we re-analyzed public datasets (MSV000085944, MSV000084008, and MSV000082493) from two healthy cohorts receiving specific drugs followed by intensive time-series sampling of multiple biofluids. The first cohort received a single dose of oral diphenhydramine (50 mg) with collection of plasma samples and skin swabs prior to (0 hour) and 24 hours after administration (0.5, 1, 1.5, 2, 4, 6, 8, 10, 12, and 24 hours).[48] The second cohort received a cocktail of orally administered probe drugs including caffeine (2 mg/kg), midazolam (0.075 mg/kg), and omeprazole (40 mg) at day 1, followed by orally administered cefprozil (500 mg) twice daily at day 2-8. Plasma samples collected at day 1 (prior to drug administration and after 5 min, 30 min, 1, 2, 4, 5, 6, and 8 hours) and fecal samples collected at day 0-9 were included in the analysis.[50] The peak list files (.mzML) were searched against the GNPS Drug Library using the library search workflow on GNPS2, with precursor and fragment ion mass tolerances of 0.02 Da, a cosine score threshold of 0.7, and minimum 2 matched peaks.

Annotations with cosine score lower than 0.9 and matched peaks fewer than 5 were removed in downstream analysis. Annotations of the drug, drug metabolites, and drug analogs were grouped based on the parent drug. The GNPS job links for are available at: https://gnps2.org/status?task=b346a5c4b70b45bebeb552e759c2bff7 , https://gnps2.org/status?task=fb2b05416ed549ad8915a8b0233d76f6 , and https://gnps2.org/status?task=2b83ce8dcae44dae8bad958e6db5765f.

***Reanalysis of public data in ReDU***

The MGF spectra files of the GNPS Drug Library were imported into the fastMASST pipeline[42,43] and searched against the GNPS/MassIVE repository using the following parameters: 0.05 for both the precursor *m/z* tolerance and fragment *m/z* tolerance, 0.8 cosine threshold, 6 minimum matched peaks, and analog search off. In order to have an overview of the drug exposures among different disease groups, the fastMASST results were merged with ReDU ("Reanalysis of Data User Interface") controlled vocabulary metadata, which is an interface linking metadata with curated controlled vocabularies to public untargeted metabolomics data files.[51] This merged table was then filtered to contain only matches to human datasets ("9606|Homo sapiens" in the NCBITaxonomy column). After filtering, the unique combinations of disease status and body parts were extracted based on the "DOIDCommonName" and the "UBERONBodyPartName" column. Detected drugs were connected to their therapeutic areas using the GNPS Drug Library metadata, separating the "infectious disease" drugs into antibiotics, antifungals, and antivirals. The heatmap was generated to visualize exposure profiles based on disease status by calculating the numbers of samples with drugs in a certain therapeutic area normalized to the numbers of samples in each sample type with at least one drug detection. The heatmap was generated using the 'ComplexHeatmap' package in R (v 2.20.0) with script provided in the code availability section.

***Reanalysis of the American Gut Project data***

In order to overview drug exposure profiles among ages and genders, the mzML files from the public metabolomics dataset of the American Gut Project (MSV000080673) were re-processed with feature extraction in MZmine 3.[83] The data were originally acquired with LC-qTOF-MS/MS and the following MZmine settings were used: For mass detection, the noise levels were 1E3 for MS1 detection and 5E1 for MS2 detection. For chromatogram building, the mass tolerance was set as 0.0050 *m/z* or 30 ppm, the minimum consecutive scans as 5, and the minimum height as 3E3. For local minimum search for chromatographic deconvolution, minimum search range was 0.1 min, minimum ratio of peak top to side was 2, and maximum peak duration was 2.0 min. The peaks were de-isotoped within 3 ppm *m/z* and 0.08 minutes retention time tolerances, then aligned with 0.0050 *m/z* or 30 ppm mass tolerance and 0.3 minutes retention time tolerance. The feature list was filtered for minimum detections in 2 samples and exported as a feature quantification table (.csv) and an MGF spectra file. The feature quantification table was subsequently filtered with R scripts to remove features with average peak areas in samples lower than 3-fold of those in blanks. For the remaining features, peak areas in samples were considered detected only if they were higher than 3-fold of the average peak areas in blanks.

Annotations were performed with the GNPS Drug Library using the Feature-Based Molecular Networking (FBMN) workflow on GNPS.[84] The spectra were filtered by removing MS/MS fragments within ±17 Da of the precursor *m/z* and to only keep the six most intense fragments in ±50 Da window. The spectra were searched against the GNPS Drug Library with precursor and fragment ion mass tolerances of 0.01 Da, a cosine score threshold of 0.7 and minimum 2 matched peaks. The GNPS job is available at: https://gnps.ucsd.edu/ProteoSAFe/status.jsp?task=649a6b4bfc5a4b1d9131b3ff23c1ae12. Annotations with cosine score lower than 0.9 and matched peaks fewer than 5 were removed in downstream analysis. The remaining annotations were manually inspected and were accepted only if two or more major fragment ions were matched.

Demographic information of the samples were collected from ReDU controlled vocabulary metadata, including "AgeInYears" column for age, "BiologicalSex" column for sex, "Country" for country of residence, and "LatitudeandLongitude" for geographic coordinates. Samples from the intensive care unit (ICU) microbiome pilot (Qiita study 2136)[85] were excluded from the analysis because their drug exposure does not resemble the general population (n = 93). In cases where individuals had more than 1 sample (n = 47), only the first collected sample was included in the analysis. Only individuals from the US, Europe, and Australia were included in the analysis, due to few samples from other regions (n = 50). Annotations were filtered based on the exposure source to exclude drugs with endogenous or diet-derived sources. Drug annotations were further grouped based on unique parent compound names, and the number of drugs detected per individual were counted to create the world map visualization (**Figure S4b**). Geographic coordinates were available for n = 1,903 individuals. The world map was generated using the "rworldmap" package in R with scripts provided in the code availability section. For the visualization based on pharmacologic classes (**Figure 1g-h**), drug annotations were further grouped based on the pharmacologic class metadata provided in the GNPS Drug Library. Each pharmacologic class was counted only once per individual when multiple drugs under the same class were detected. Age groups were defined in 10-years intervals (nine intervals for age 0-90 years old), and individuals with no information on age and sex (n = 148) and individuals >90 years (n = 2) were excluded from the analysis. To adjust for the varying number of samples across age and sex, normalization was performed by dividing the number of observations by the total number of samples within each age group and sex.

### *Extraction and LC-MS/MS profiling of human feces with HIV infection*

*Sample extraction.* The stool samples were prepared with a recently developed automatic pipeline for simultaneous metagenomics-metabolomics extractions.[86] Specifically for the metabolite extraction, swabs were added into Matrix Tubes (ThermoFisher Scientific, MA, USA) containing 400 µL of 95% (v/v) ethanol and capped with the automated instrument Capit-All (ThermoFisher Scientific, MA, USA). The tubes were shaken for 2 minutes (1,200 rpm, the SpexMiniG plate shaker) followed by centrifugation for 5 minutes (2,700 g). The supernatant was then transferred to a deep well plate using an 8-channel pipette, concentrated to dryness with a vacuum centrifuge concentrator (room temperature; ~5 hours) and stored at -80 °C. Prior to instrumental analysis, the plates were redissolved (10 minute sonication) in 200 µL of 50% acetonitrile (v/v) with 100 µg/L sulfadimethoxine as the internal standard. Plates were centrifuged at 450 g for 10 minutes, and 150 µL supernatants were collected for instrumental analysis.

*Instrumental analysis.* The fecal extracts were injected (5 µL) into a Vanquish ultra-high-performance liquid chromatography (UHPLC) system coupled to a Q Exactive quadrupole orbitrap mass spectrometer (Thermo Fisher Scientific, Waltham, MA). A Kinetex polar C18 column (150 × 2.1 mm2, 2.6 µm particle size, 100 A pore size; Phenomenex, Torrance) was employed with a SecurityGuard C18 column (2.1 mm ID) at 30 °C column temperature. The mobile phases (0.5 mL/min) were 0.1% formic acid in both water (A) and ACN (B) with the following gradient: 0-1 min 5 % B, 1-7 min 5-99 % B, 7-8 min 99 % B, 8-8.5 min 99-5% B, 8-10 min 5% B. The mass spectrometer was operated in positive heated electrospray ionization with the following parameters: sheath gas flow, 53 AU; auxiliary gas flow, 14 AU; sweep gas flow, 3 AU; auxiliary gas temperature, 400 °C; spray voltage, 3.5 kV; inlet capillary temperature, 269 °C; S-lens level, 50 V. MS1 scan was performed at *m/z* 100-1500 with the following parameters: resolution, 35,000 at *m/z* 200; maximum ion injection time, 100 ms; automatic gain control (AGC) target, 5.0E4. Up to 5 MS/MS spectra per MS1 scan were recorded under the data-dependent mode with the following parameters: resolution, 17,500 at *m/z* 200; maximum ion injection time, 100 ms; AGC target, 5.0E4; MS/MS precursor isolation window, *m/z* 3; isolation offset, *m/z* 0.5; normalized collision energy, a stepwise increase from 20 to 30 to 40 %; minimum AGC for MS/MS spectrum, 5.0E3; apex trigger, 2 to 15 s; dynamic precursor exclusion, 10 s.

*Feature extraction and data analysis.* The raw spectra were converted to mzML files using MSconvert (ProteoWizard)[87] followed by feature extraction using MZmine 4[83] with the following parameters: For mass detection, the noise factor was 3.5 for MS1 and 2.5 for MS2. For chromatogram building, the mass tolerance was set as 0.01 *m/z* or 10 ppm, the minimum consecutive scans as 3, and the minimum height as 5E2. Chromatograms were smoothed with the Savitzky Golay algorithm, followed by local minimum search for chromatographic deconvolution with minimum search range of 0.2 min, minimum ratio of peak top to side of 2, and maximum peak duration of 0.5 min. The peaks were de-isotoped within 5 ppm *m/z* and 0.05 minutes retention time tolerances, aligned with 0.01 *m/z* or 20 ppm mass tolerance and 0.3 minutes retention time tolerance, then gap-filled with 20 ppm *m/z* tolerance and 0.3 minutes retention time tolerance. The feature list was blank-subtracted with 300% fold change increases compared with maximum blank values and 30 minimum number of detection in blanks. The feature list was then exported as a feature quantification table (.csv) and an MGF spectra file and used without post-filtering.

The features were annotated for drugs and drug analogs using the GNPS Drug Library with the same procedure as for the American Gut Project data. The GNPS job is available at: https://gnps.ucsd.edu/ProteoSAFe/status.jsp?task=90b6edcfb8fc43dab54d2d7951a3291a. Peak areas of annotated drugs lower than 1E4 were replaced by zero to avoid false detections caused by instrument noises. Representative drug annotations, including all annotated ARVs and drugs used in the microbial incubations, were validated by analytical standards (**Figure S8**).The features were annotated for *N*-acyl lipids using a recently developed a spectral library for 851 *N*-acyl lipids[70] with the following parameters for the FBMN workflow: 0.02 Da precursor and fragment ion mass tolerance and library search with a cosine score threshold of 0.7 and minimum 4 matched peaks. The GNPS job is available at https://gnps2.org/status?task=ee34ee95908749dd81ee9a62fbdac98e. The heatmaps to demonstrate drug annotations were generated using the 'ComplexHeatmap' package in R (v 2.20.0). People with HIV were clustered into four groups based on their ARVs profiles by hierarchical clustering under the Ward's linkage method and Euclidean distance matrix. Boxplots representing the peak areas of *N*-acyl lipids in different drug exposure groups were generated using the 'ggplot2' package in R (v 3.5.1), with R scripts provided in the code availability section.

***Synthetic microbial community culture for drug metabolism***

*Chemicals.* Drugs for bacterial metabolism screening were purchased from Sigma-Aldrich (St. Louis, MO, USA; see detailed list in **Table S4**). All solvents used were Optima LC-MS grade from Fisher Scientific (Pittsburgh, PA, USA). We note that quinine and quinidine were included in the drug metabolism assays because they were initially annotated based on MS/MS spectra matches (level 2/3 according to the 2007 Metabolomics Standards Initiative)[77]. After we acquired the standards, their retention times do not match the peaks in fecal samples. Therefore, quinine and quinidine were not considered in downstream analysis.

*Synthetic microbial community culture.* The 111 bacterial strains used in this study are a selection of the human gut microbiota (hCom) previously published[60] that was optimized in the Zengler lab to accomplish uniform growth (**Table S5**). A normalized working stock solution at $OD_{600}$ = 0.01 was created for the 111 bacterial strains and stored at -80 ℃ until needed. For culturing, six groups were created and include the full hCom, the hCom-reduced (*Coprococcus comes* and *Coprococcus eutactus* were omitted due to their dominance in the community), fast growers, medium-fast growers, medium-slow growers, and slow growers (**Table S5**). To create the hCom and hCom-reduced communities, the normalized stock of a bacteria was thawed, and a specific volume was taken and combined to generate the groups. To create the hCom-reduced, each bacteria was added based on their groups (fast-growers, 0.5 µL; medium-fast growers, 5 µL; medium-slow, 50 µL (not including the two species of *Coprococcus*); and slow growers, 500 µL). *Coprococcus comes* and *Coprococcus eutactus* were added at 0.5 µL to finalize the assembly of the full hCom. To create the other group communities, 500 µL of each bacteria of each group were combined (**Table S5**). The groups were then diluted 1/100 at a final $OD_{600}$ of 0.0001 and cultured anaerobically in BHI medium (**Table S6**) at 37 ℃ in an anaerobic chamber (10% $CO_2$, 7.5% $H_2$, 82.5% $N_2$).

*Drug metabolism assays.* A 10 mM stock solution was created for each drug and diluted to 200 µM with $H_2O$. Drugs were combined to create four different mixtures for the screening assay (mix1: sertraline, quinidine, venlafaxine; mix2: cetirizine, losartan, quetiapine, ritonavir; mix3: darunavir, quinine, atorvastatin, omeprazole; mix4: loratadine, terbinafine, trazodone, abacavir). Each drug was added to a final concentration of 2 µM (2 µL spike) after 24 h of bacterial acclimation. Cultures were immediately extracted after drug addition as the 0 h samples. Cultures from the fast and medium-fast groups were extracted after 48 h of growth and the cultures from the remaining groups were stopped at 72 h. Medium controls were included by adding drug mixtures to the BHI medium without microbial communities and were included at both 0 h and 72 h. All conditions were performed in triplicates.

*Extraction of drugs and metabolites for LC-MS analysis.* Bacterial cultures were extracted using 600 µL of pre-chilled 50% (v/v) MeOH, followed by an overnight incubation at 4℃. Samples were dried using a CentriVap and stored at -80℃ until LC-MS/MS analysis. Samples were resuspended in 200 µL of pre-chilled 50% (v/v) MeOH with 100 µg/L sulfadimethoxine as an internal standard, incubated at -20℃ overnight, centrifuged at 15,000 x g for 10 min, and 150 µL of the supernatant was transferred to 96 shallow well-plate for LC-MS analysis. Samples were analyzed with the same LC-MS/MS method for the HIV feces samples, except the following parameters: additional LC washing gradient after sample analysis, 8.5-9 min 5% B, 9-9.5 min 5-99% B, 9.5-10.5 min 99% B, 10.5-11 min 99-5% B, 11-12.5 min 5% B; MS1 scan range, *m/z* 100-100; AGC target, 1E6; dd-MS2 AGC target, 5.0E5; MS/MS precursor isolation window, *m/z* 1.

The raw spectra were converted to mzML files followed by feature extraction using MZmine 4 with the following parameters: For mass detection, the noise factor was 3.0 for MS1 and 2.0 for MS2. For chromatogram building, the mass tolerance was set as 0.002 *m/z* or 10 ppm, the minimum consecutive scans as 4, and the minimum height as 1.5E5. Chromatograms were smoothed with the Savitzky Golay algorithm, followed by local minimum search for chromatographic deconvolution with minimum search range of 0.05 min, minimum ratio of peak top to side of 2, and maximum peak duration of 1.5 min. The peaks were de-isotoped within 3 ppm *m/z* and 0.04 minutes retention time tolerances, aligned with 0.002 *m/z* or 10 ppm mass tolerance and 0.07 minutes retention time tolerance, then gap-filled with 10 ppm *m/z* tolerance and 0.05 minutes retention time tolerance.

The features were annotated for drugs and drug analogs using the GNPS Drug Library with the same procedure as for the American Gut Project data. The GNPS job is available at: https://gnps2.org/status?task=128c85f310844f2d90d71bcf1c107afb.

*Co-migration analysis with fecal extracts.* Four samples from the bacterial cultures were analyzed again with selected fecal samples from the HIV infection cohort to confirm the retention time and MS/MS spectral matching of the drug analogs. One medium-slow grower at 72 hours was used to confirm the ritonavir analog at *m/z* 737.31, one medium-fast grower at 72 hours for the atorvastatin analog at *m/z* 573.28, one medium-fast grower at 72 hours for abacavir analogs at *m/z* 247.13 and 319.15, and one hCom-reduced culture at 0 hour for the omeprazole analog at *m/z* 330.13. Fecal samples with the highest analog intensities were used. Extracts of the bacterial cultures or feces were first diluted to similar concentrations with 50% (v/v) methanol based on peak areas of the drug analogs in the initial sample analysis, and then were mixed at 1:1 volume ratio. The selected bacterial culture samples, fecal samples, and mixture samples were co-analyzed using the same LC-MS/MS method for the bacterial cultures.

***Statistical Analysis.***

A chi-square test was performed to compare the number of drugs detected in fecal samples from different countries (**Figure S4b**). Kruskal-Wallis tests followed by a pairwise Wilcoxon test were performed with Benjamini-Hochberg correction to compare the levels of *N*-acyl lipids in the four drug exposure groups (**Figure 2d**, **Figure S7b**). Adjusted p-values $<0.05$ were considered as statistically significant. Pairwise Wilcoxon tests were performed to compare the drug analog intensities in bacterial cultures at 0 hour and 72 hours (**Figure S6**). Due to the small sample size ($n = 3$ in each group), p-values $<0.1$ were considered as statistically significant.
