## Supplementary Figures for "Empirically establishing drug exposure records directly from untargeted metabolomics data"

[19] Department of Medicinal Chemistry, University of Washington, Seattle, WA, USA
[20] Current address: Biological Sciences Division, Pacific Northwest National Laboratory, Richland, WA, USA
[21] Department of Computer Science and Engineering, University of California Riverside, Riverside, CA, USA
[22] Center for Microbiome Innovation, University of California San Diego, La Jolla, CA, USA.
[23] Department of Computing Science, University of Alberta, Edmonton, AB T6G 2E8, Canada
[24] Department of Psychiatry and Behavioral Sciences, Duke University, Durham, NC, 27708, USA
[25] Duke Institute of Brain Sciences, Duke University, Durham, NC, USA
[26] Department of Medicine, Duke University, Durham, NC, USA
[27] Chiba University, UC San Diego Center for Mucosal Immunology, Allergy, and Vaccines (CU-UCSD cMAV), La Jolla, CA, USA
[28] Program in Materials Science and Engineering, University of California, San Diego, 9500 Gilman Drive, La Jolla, CA 92093-0418, USA
[29] Department of Computer Science and Engineering, University of California San Diego, La Jolla, CA, USA
[30] Shu Chien-Gene Lay Department of Bioengineering, University of California San Diego, La Jolla, CA, USA
[31] Halıcıoğlu Data Science Institute, University of California San Diego, La Jolla, CA, USA
† Haoqi Nina Zhao, Kine Eide Kvitne, and Corinna Brungs contributed equally to this work.
* Author to whom correspondence should be addressed.

**Supplementary Text 1. Detailed steps for the collection of MS/MS reference spectra for drugs using the MS$^n$ Python script.**

Reference spectra of drugs and known drug metabolites were collected from the GNPS spectral library and the MS$^n$ library (generated under MZmine version 3.4.0) using a Python script developed in the MS$^n$Lib workflow.[26] Detailed steps include:

- PubChem database search to find missing or wrong structural information
    - Query by PubChem CID, CAS, or name
- Structure cleanup and standardization
    - Computing the canonical, isomeric SMILES, InChI, InChIKey, and first block of InChIKey to remove stereochemistry
- Pubchem database search to get all synonyms
    - Query by structural information, including InChIKey , SMILES, or InChI
- UniChem database search to extract database identifiers, e.g., Drugbank ID, ChEMBL ID
    - Query based on InChIKey
- Extract missing database identifiers from the PubChem synonyms column
- Broad Institute Drug Repurposing Hub database search for drug information
    - Query based on first block of InChIKey
    - Output: Preclinical, Phase 1, Phase 2, Phase 3, Launched, or Withdrawn
- ChEMBL database search for drug information
    - Query based on ChEMBL ID or InChIKey
    - Output: clinical phase as -1, 0, 0.5, 1, 2, 3, 4
- DrugBank database search for drug information
    - Query based on Drugbank ID, InChIKey, PubChem CID, ChEMBL ID, UNII ID, CAS, first block of InChIKey, or compound name
    - Output: approved, withdrawn, vet_approved, nutraceutical, investigational, illicit, experimental
    - Approved and withdrawn translated to clinical phase 4
- DrugCentral database search for drug information
    - Query based on first block of InChIKey
    - Output: approval agency, e.g., EMA, FDA, KFDA (Korean Food and Drug Administration), or PMDA (Pharmaceuticals and Medical Devices Agency, Japan)
    - Approval by agency converted to clinical phase 4
- Clinical phase description for each database converted to number (0, 0.5, 1, 2, 3, 4)
- Get the highest clinical phase for each compound based on all gathered information from the databases

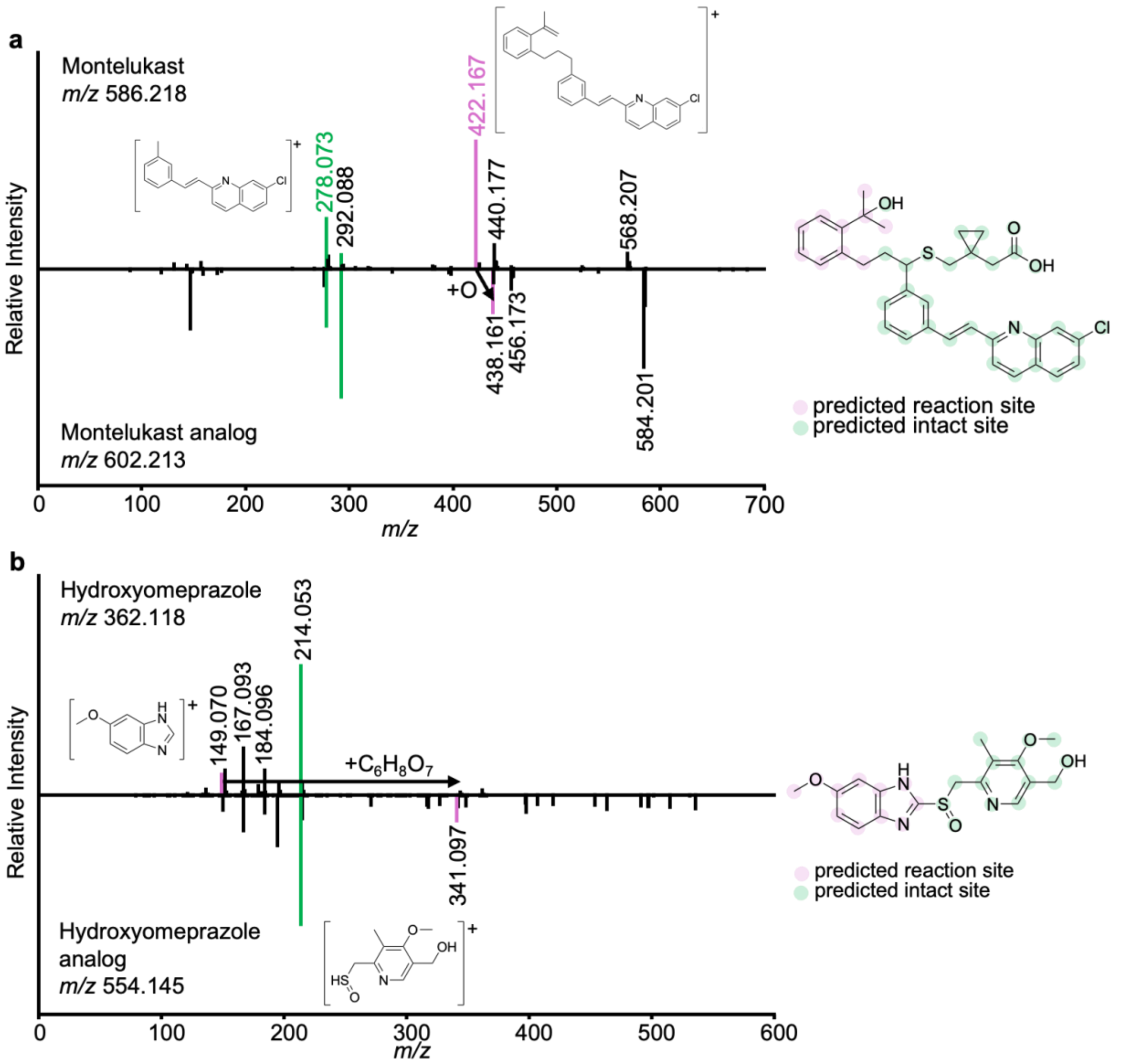


**Figure S1. Additional examples of structural modification sites of the drug analogs predicted by ModiFinder.**[32] **a,** Montelukast analog with a delta mass of +15.99 Da (+O). Matched ions (*m/z* 278.073, 292.088) between the drug and the analog suggested unmodified substructures on the drugs. Shifted ions with 15.99 Da mass difference (*m/z* 422.167 to 438.161, 440.177 to 456.173, 568.207 to 584.201) suggested modification sites. **b,** Hydroxyomeprazole analog with a delta mass of +192.03 Da (+$C_6H_8O_7$). Matched ions: *m/z* 167.093, 184.096, 214.053; shifted ions: *m/z* 149.070 to 341.097. Pink traces highlight modified spectra and substructures, while green traces highlight unmodified ones.

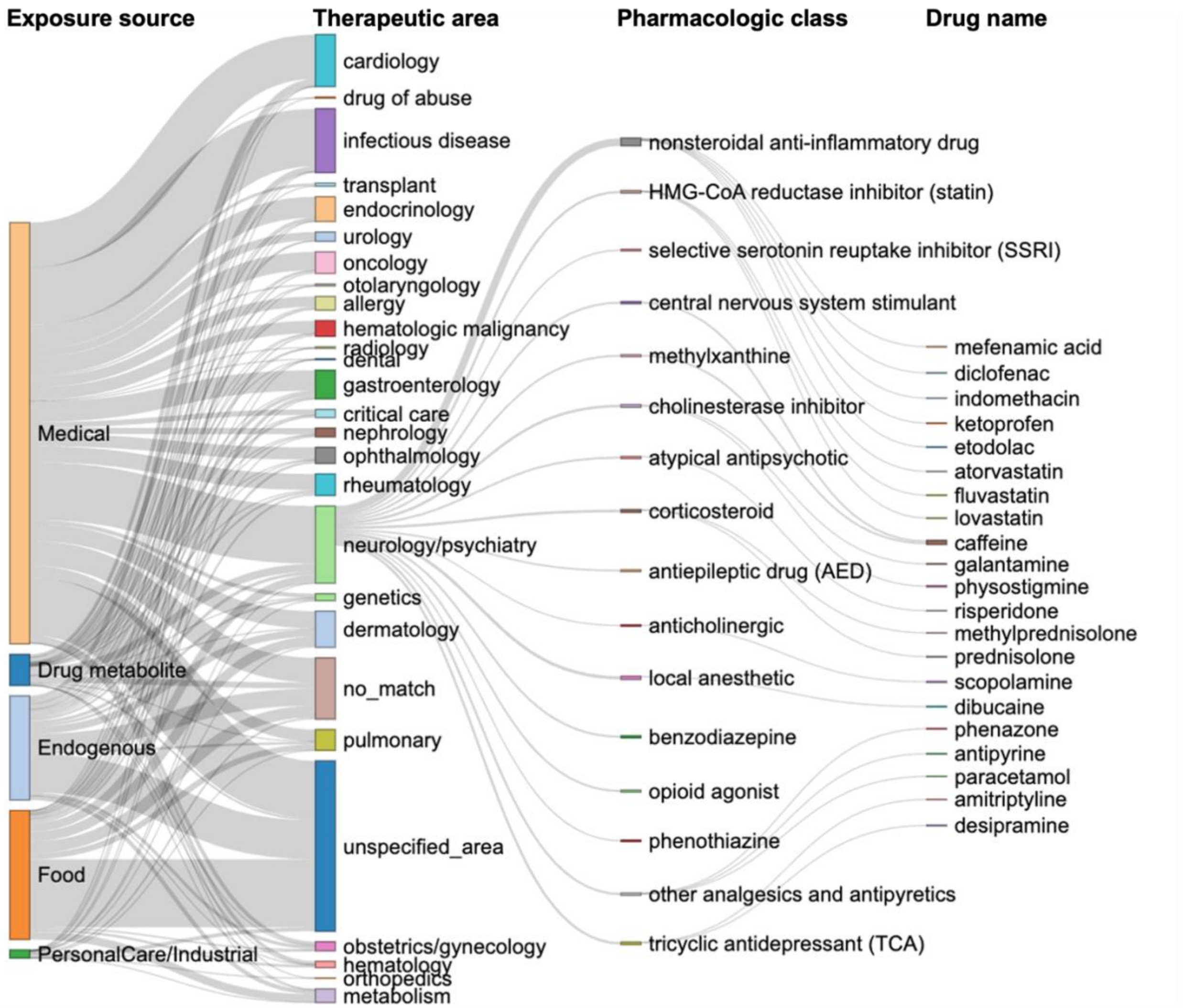


**Figure S2.** Overview of the ontology-based drug metadata based on the numbers of reference spectra, highlighting common therapeutic areas, pharmacologic classes, and specific drugs in the neurology/psychiatry category. The width of the bars and lines reflect the number of reference spectra in each category.

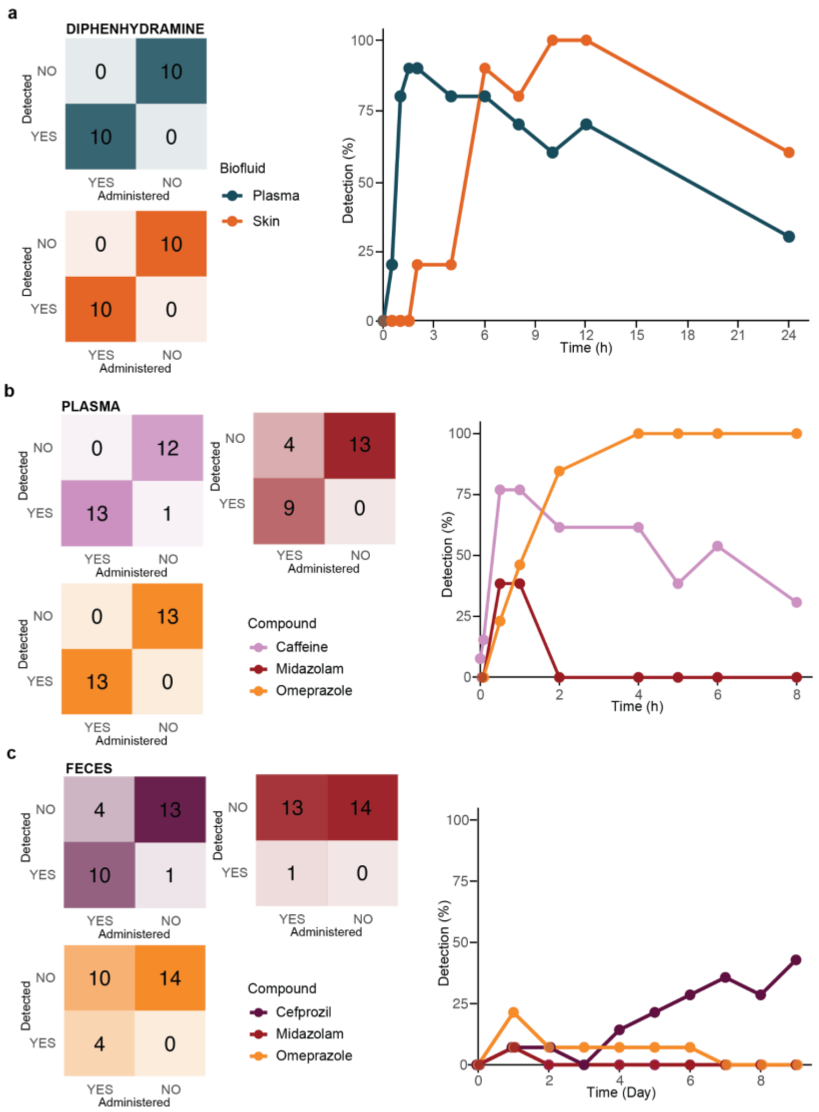


**Figure S3. Empirical drug readout in healthy individuals known to be receiving specific drugs in pharmacokinetic studies.**[48,50] **a,** Detection frequencies of diphenhydramine in plasma and skin samples from 10 individuals receiving a single dose of oral diphenhydramine (50 mg). The confusion matrix summarizes drug detections across all time points, while the line chart demonstrates detections before (0 hour) and 24 hours (0.5, 1, 1.5, 2, 4, 6, 8, 10, 12, and 24 hours) after drug administration. Samples collected before drug administration were used as "Administered-NO" in the confusion matrix. **b,** Detection frequencies of the administered drugs in plasma samples from 13 individuals receiving a cocktail of oral probe drugs, including caffeine (2 mg/kg), midazolam (0.075 mg/kg) and omeprazole (40 mg). Plasma samples were collected prior to and 5 min, 30 min, 1, 2, 4, 5, 6, and 8 hours after administration of the drug cocktail. **c,** Detection frequencies of drugs in fecal samples from 14 individuals receiving the drug cocktail on day 1, followed by cefprozil from day 2-8 (500 mg twice daily). The confusion matrices and line charts in panel b-c were constructed in the same way as panel a. Caffeine was not detected in the fecal samples and therefore not included in panel c.

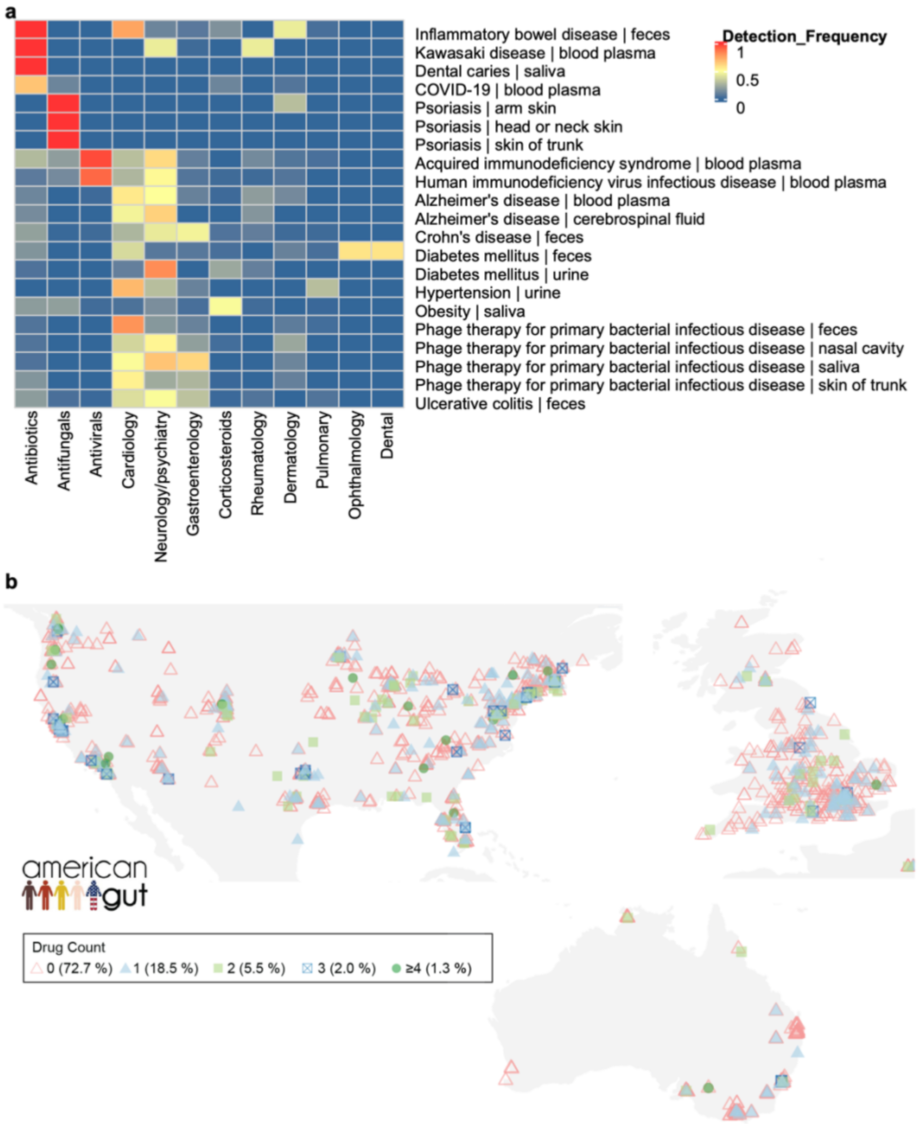


**Figure S4. Drug exposure profiles among different disease types and geolocations by re-analyzing public metabolomics data. a,** The GNPS Drug Library was searched against ReDU[50] samples with curated disease ontology (n = 1,773) using fastMASST.[42,43] Rows of the heatmap represent unique combinations of the disease types and sample types, and columns represent drug categories based on disease areas. The heatmap was colored by the drug detection frequencies, defined as the number of observations normalized to the number of samples in each sample type with at least one drug detection. **b,** Number of drugs detected in each individual in the American Gut Project[33] visualized on a world map. The number of drugs used by each individual from the US (n = 1245, 0 drug; 68.8%, 1 drug; 20.4%, 2 drugs; 6.4%, 3 drugs; 2.6%, ≥4 drugs; 1.8%), Europe (n = 533, 0 drug; 79.9%, 1 drug; 15.2%, 2 drugs; 3.9%, 3 drugs; 0.75%, ≥4 drugs; 0.19%), and Australia (n = 125, 0 drug; 80.0%, 1 drug; 14.4%, 2 drugs; 3.2%, 3 drugs; 1.6%, ≥4 drugs; 0.8%) varied significantly (chi-square test; $\chi^2$ (8, n = 1,903) = 33, $p = 5.3 \times 10^{-5}$). Only samples from the UK (n = 488) are visualized in the map due to few samples (n = 45) from the rest of Europe.

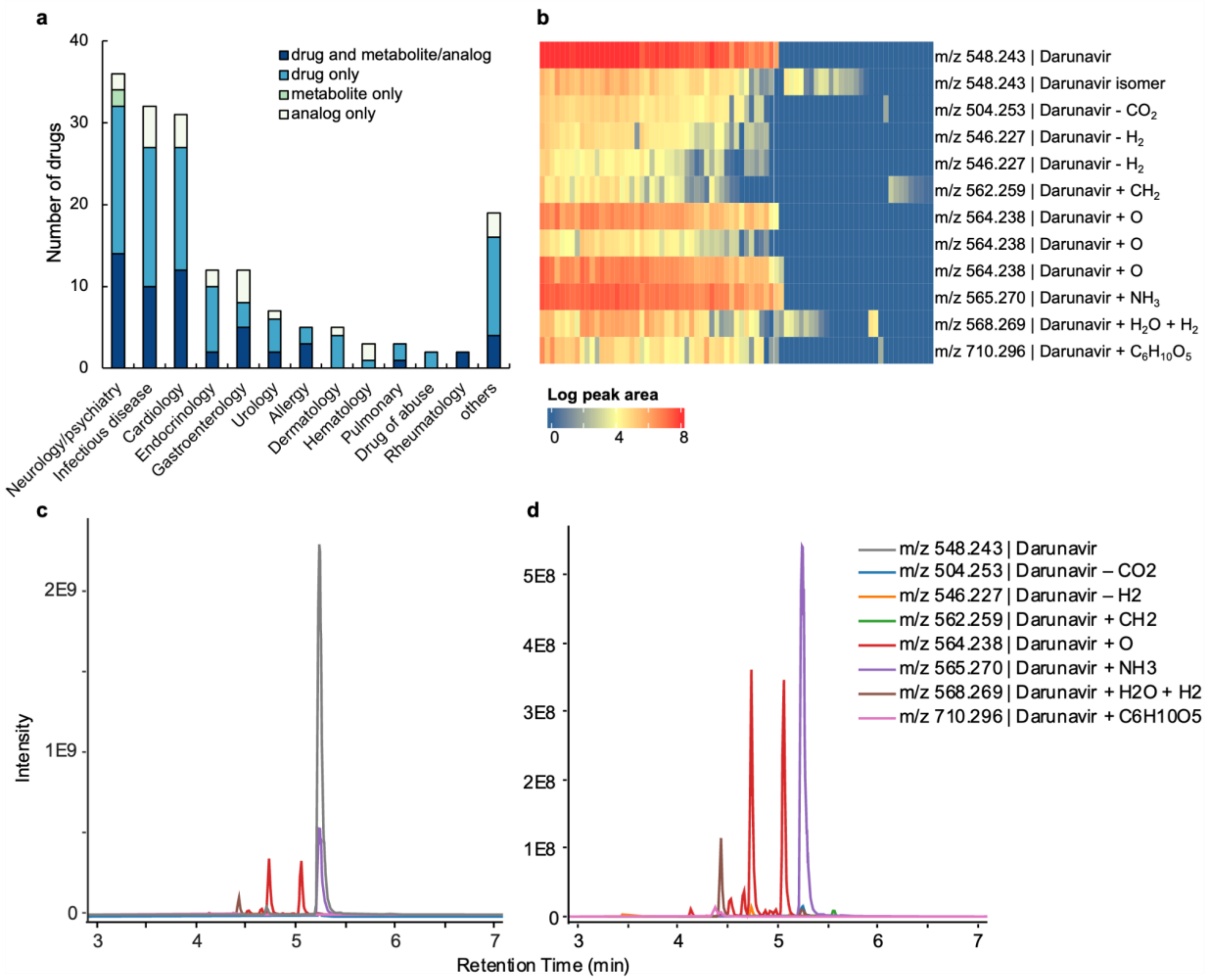


**Figure S5. Drug analog annotations in fecal samples from the HIV Neurobehavioral Research Center (HNRC) cohort (n = 322) of people with (n = 222) and without HIV (n = 100). a,** Number of drugs detected in each disease area, colored based on detection types as the parent drug only, drug with metabolites/analogs, or only drug analogs. **b,** Peak area visualization of darunavir and all darunavir analogs. Each column represents one sample and each row represents one annotation. Samples without darunavir detections are not shown. The atomic changes of the drug analogs were based on [M+H]+ ion of the parent drug. **c,** Chromatograms of darunavir and all analogs. Two analogs (*m/z* 565.27, 504.25) are identified as darunavir in-source fragments due to highly correlated retention times and peak shape. Other analogs are likely darunavir metabolites, transformation products, or isomers. **d,** Chromatograms of all darunavir analogs without the darunavir ion to enhance visualization. The atomic changes of the drug analogs were based on [M+H]+ ion of the parent drug.

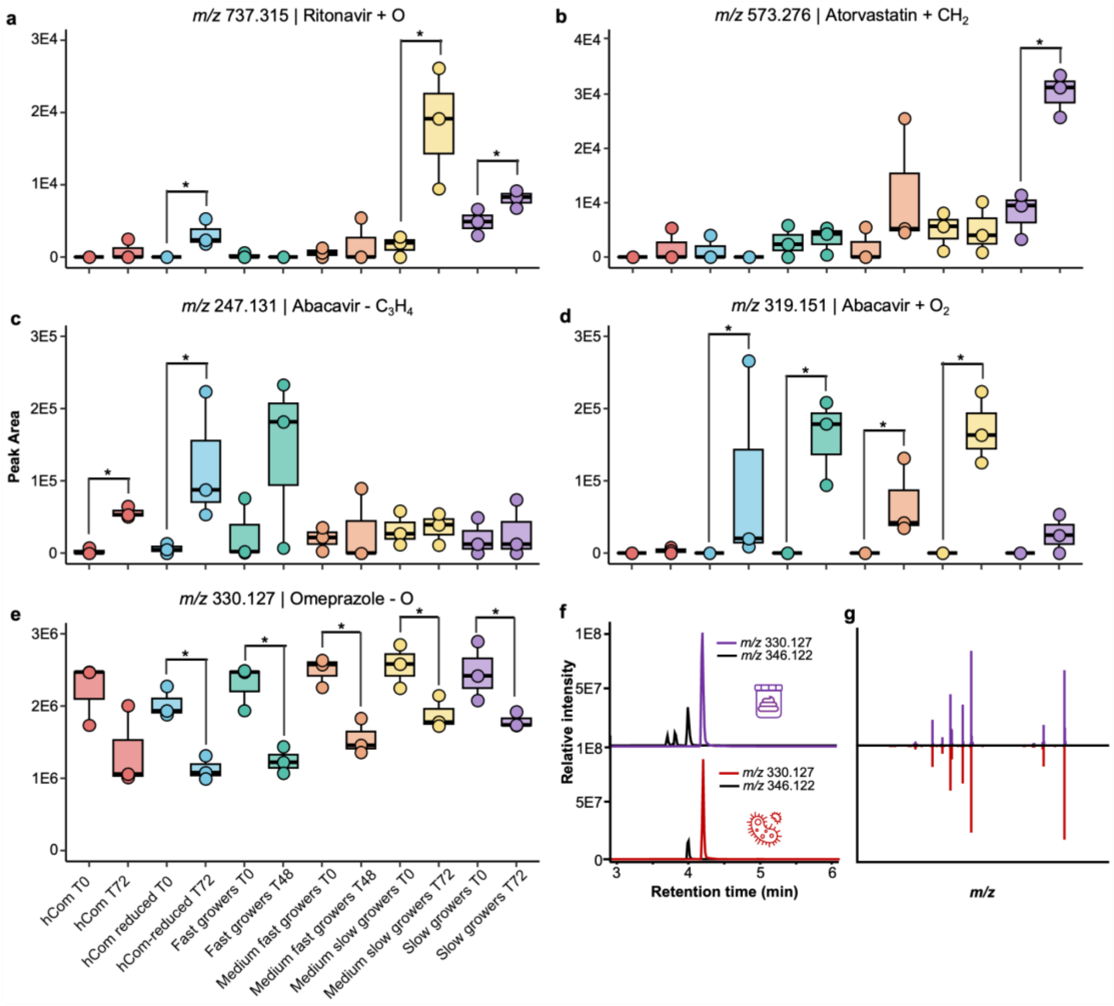


**Figure S6. Drug analogs observed in human fecal samples can be produced by microbial metabolism. a-e,** Peak area variation for 5 drug analogs at 0 and 72 h of microbial incubation, including ritonavir analog with delta mass of +15.99 Da (*m/z* 737.315; **a**), atorvastatin analog with delta mass of +14.02 Da (*m/z* 573.276; **b**), abacavir analog with delta mass of -40.03 Da (*m/z* 247.131; **c**), abacavir analog with delta mass of +31.99 Da (*m/z* 319.151; panel **d**), and omeprazole analog with delta mass of -15.99 Da (*m/z* 330.127; **e**). Panels a-d share the same x-axis labels as panel e. The atomic changes of the drug analogs were based on $[M+H]^+$ ion of the parent drug. Drugs were cultured with six groups of synthetic microbial community, including the full hCom, the hCom-reduced (*Coprococcus comes* and *Coprococcus eutactus* omitted due to their dominance in the community), fast growers, medium-fast growers, medium-slow growers, and slow growers. Non-parametric Wilcoxon tests were performed to compare peak areas at 0 h and 72 h, and p-values < 0.1 were marked with asterisks. **f-g,** Retention time (**f**) and MS/MS spectra mirror matches (**g**) for the omeprazole analog in human fecal samples and the microbial incubations. Purple traces represent the fecal samples, while red traces represent the microbial incubation. The extracted ion chromatogram of the parent drug (*m/z* 346.122) is additionally visualized in black for peak area comparison.

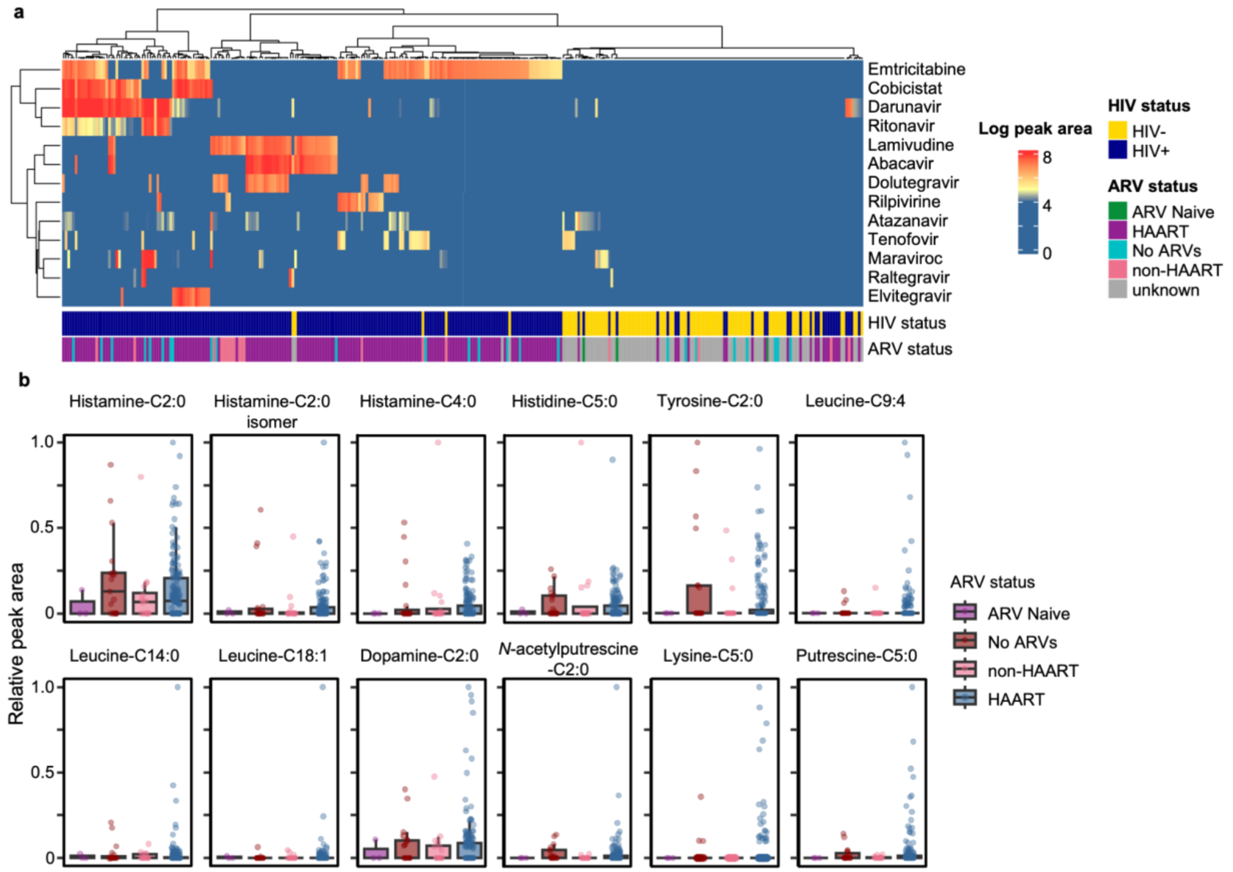


**Figure S7. Comparison of sample clustering based on empirical drug records from the GNPS Drug Library and clinical metadata. a,** Peak area visualization of antiretrovirals (ARVs) detected in the HIV Neurobehavioral Research Center cohort (n = 322 fecal samples). Each column represents one sample and each row represents one ART. Peak areas of drug, drug metabolite, and drug analogs belonging to the same parent drug are added and log-transformed. Rows and columns of the heatmap were arranged by hierarchical clustering analysis with Ward's linkage and Euclidean distance. The heatmap columns were noted with the HIV serostatus and ARV usage status from the clinical metadata. **b,** Sample-to-sample peak areas of the *N*-acyl lipids in people with HIV, separated by the ARV exposure status reported in clinical metadata. ARV-naïve, never received ARV; no ARV, no current ARV use; non-HAART, currently using less than three ARVs; HAART, currently using three or more ARVs. The peak area was normalized to the maximum value observed for the specific compound. No significant difference was observed for the *N*-acyl lipid levels based on clinical self-reported ART exposure status (non-parametric Kruskal-Wallis tests p-value > 0.05). Horizontal lines indicate the median value, the first (lower) and third (upper) quartiles are represented by the box edges, and vertical lines (whiskers) indicate the error range which is 1.5 times the interquartile range.

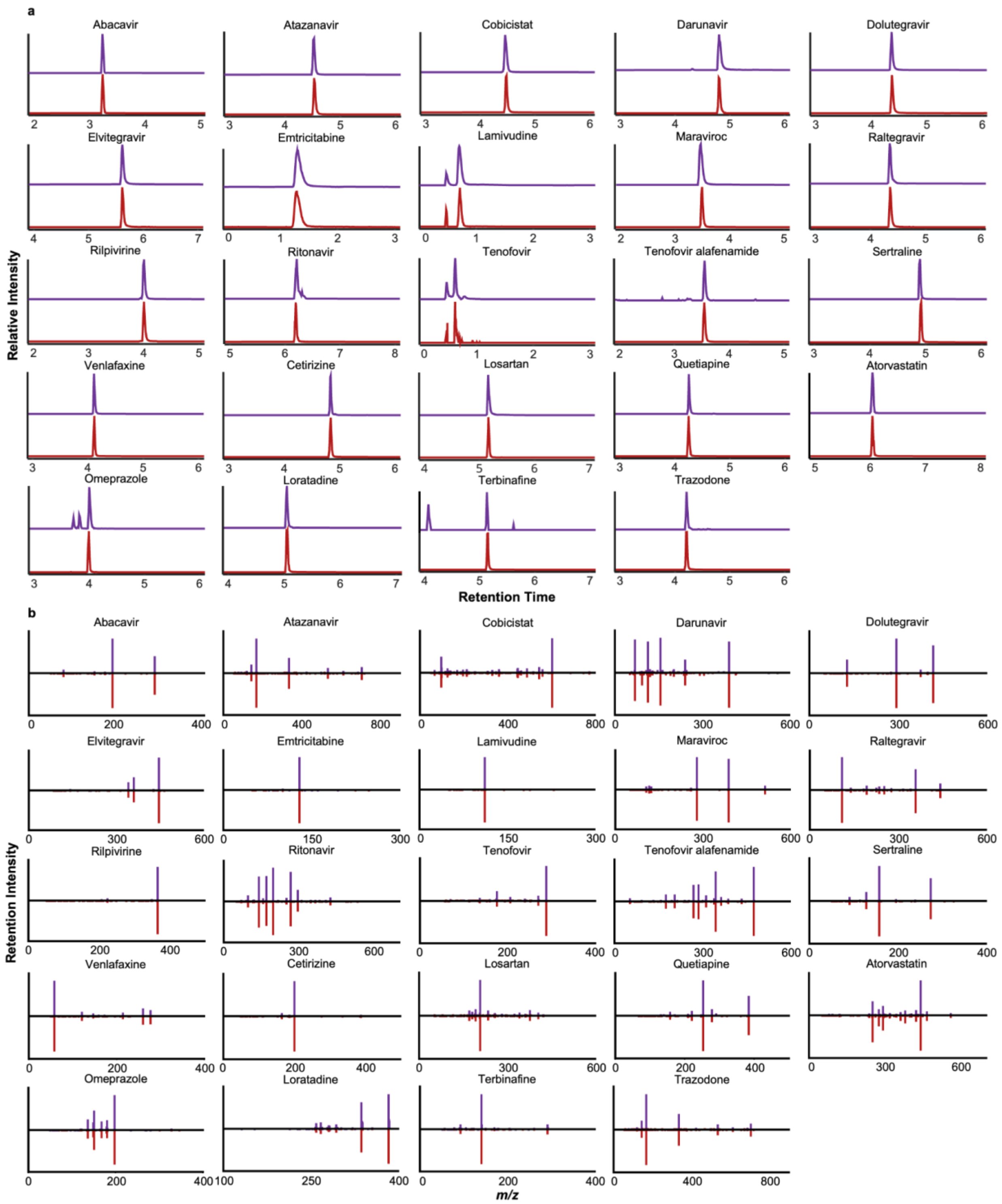


**Figure S8. (a) Retention time and (b) MS/MS spectra mirror matches for drugs observed in the HNRC cohort with analytical standards.** Purple traces represent the fecal samples, while red traces represent the analytical standards.
